# Genomic insights into the Archaea inhabiting an Australian radioactive legacy site

**DOI:** 10.1101/728089

**Authors:** Xabier Vázquez-Campos, Andrew S. Kinsela, Mark W. Bligh, Timothy E. Payne, Marc R. Wilkins, T. David Waite

## Abstract

During the 1960s, small quantities of radioactive materials were co-disposed with chemical waste at the Little Forest Legacy Site (LFLS, Sydney, Australia). The microbial function and population dynamics in a waste trench during a rainfall event have been previously investigated using shotgun metagenomics. This revealed a broad abundance of candidate and potentially undescribed taxa in this iron-rich, radionuclide-contaminated environment.

Here, applying genome-based metagenomic methods, we recovered 37 refined archaeal MAGs (≥50% completeness, ≤10% redundancy) from 10 different major lineages. They were, mainly, included in four DPANN lineages without standing nomenclature (LFWA-I to IV) and *Methanoperedenaceae* (ANME-2D).

While most of the new DPANN lineages show reduced genomes with limited central metabolism typical of other DPANN, the orders under the LFWA-III lineage, ‘*Ca.* Gugararchaeales’ and ‘*Ca.* Anstonellales’, may constitute distinct orders with a more comprehensive central metabolism and anabolic capabilities within the *Micrarchaeota* phylum.

The new *Methanoperedens* spp. MAGs, together with previously published data, suggests metal ions as the ancestral electron acceptors during the anaerobic oxidation of methane while the respiration of nitrate/nitrite via molybdopterin oxidoreductases would have been a secondary acquisition. The presence of genes for the biosynthesis of polyhydroxyalkanoates in most *Methanoperedens* also appears to be a widespread characteristic of the genus for carbon accumulation.

We formally propose 22 new candidate taxa based on data analysed in this manuscript, as well as four missing formal taxa definitions and two new candidate species based on extant data. We present evidence of four new DPANN lineages and six non-conspecific *Methanoperedens*, while exploring their uniqueness, potential role in elemental cycling, and evolutionary history.

## Introduction

The Little Forest Legacy Site (LFLS) is a low-level radioactive waste disposal site in Australia that was active between 1960 and 1968. During this period, small quantities of both short and long-lived radionuclides (including plutonium and americium) were disposed of in three-metre deep, unlined trenches, and covered with a shallow (1 m layer) of local soil, as was common practice at the time (Payne, 2012). Since its closure, the periodic occurrence of intense rainfall events leads to the filling of the former trenches with surface waters, which, depending on preceding conditions, can result in the trench water discharging into the surface and near-surface soils in a ‘bath-tub’-like mechanism (Payne et al., 2013, 2020). Aside from providing a mechanism enabling the export of contaminants from the legacy trenches (Payne et al., 2013), the periodic influx of oxic waters into these natural reducing zones results in transitory microbial population shifts associated with pronounced elemental cycling (Vázquez-Campos et al., 2017; Kinsela et al., 2021). Our previous research showed that the archaeal community in the LFLS waste trenches, while constituting a minor portion of the whole microbial community, included a number of potentially interesting members, either in terms of phylogeny and/or functionality (Vázquez-Campos et al., 2017).

The domain Archaea constitutes by far the most understudied domain of life. Archaea have been traditionally regarded as biological oddities and marginal contributors to the global geochemical elemental cycles, with most living in extreme environments (Dombrowski et al., 2019). However, the continuous development of cultivation-independent techniques in recent decades has completely changed this perception. For example, *Thaumarchaeota* Archaea are now known to be key components in the global nitrogen cycle (Kitzinger et al., 2019) and can even induce systemic resistance to pathogens in plants (Song et al., 2019). Some *Euryarchaeota*, which are capable of the anaerobic oxidation of methane (AOM), consume up to 80–90% of the methane produced in environments such as rice fields and ocean floors before this highly deleterious greenhouse gas can be released into the atmosphere (Reeburgh, 2007).

The DPANN Archaea, so-named from its constituent lineages when it was defined (*Diapherotrites*, *Parvarchaeota*, *Aenigmarchaeota*, *Nanohaloarchaeota* and *Nanoarchaeota*) (Rinke et al., 2013), is an archaeal superphylum generally characterised by the small size of its members (excluding *Altiarchaeota*). Their diminutiveness not only relates to their reduced physical size, many of which are capable of passing through a standard 0.2 µm filter (Luef et al., 2015), but also due to their small genome sizes, i.e. many are below 1 Mbp and with an estimated maxima of ∼1.5 Mbp. Many of the DPANN with extremely reduced genomes lack biosynthetic pathways for amino acids, nucleotides, vitamins, and cofactors, which aligns with a host-dependent lifestyle. Others, towards the higher end of the genome size scale, typically encode for some or most of the aforementioned pathways and are regarded as free-living. Irrespective of size, only a limited number of organisms from this lineage have been studied in detail (Waters et al., 2003; Baker et al., 2006; Castelle et al., 2015; Youssef et al., 2015; Golyshina et al., 2017).

Another Archaea taxon that has attracted substantial attention during the last few years are the anaerobic methane oxidisers ANME-2d (Mills et al., 2003) or AAA (AOM-associated Archaea) (Knittel and Boetius, 2009), now referred to as ‘*Ca.* Methanoperedens’ (simply *Methanoperedens* henceforth). These archaea have been purported to couple the oxidation of methane, at its source, with the reduction of nitrate (Haroon et al., 2013), nitrite (Arshad et al., 2015), Fe(III) (Ettwig et al., 2016; Cai et al., 2018), Mn(IV) (Ettwig et al., 2016) or even Cr(VI) (Lu et al., 2016), constituting an important environmental pathway in mitigating the release of methane into the atmosphere upon a global scale.

Here we describe the results from the analysis of 37 new archaeal MAGs with different degrees of completeness, derived from samples collected in a legacy radioactive waste trench at the LFLS during a redox cycling event. We formally propose 22 new candidate taxa based on data analysed in this manuscript, as well as four missing formal taxa definitions and two new candidate species based on extant data. We present evidence of four new DPANN lineages and six non-conspecific *Methanoperedens* sp. genomes, while exploring their uniqueness and potential role in elemental cycling.

## Material and Methods

### Recovery and assessment of community genomes

QC and pre-processing of raw sequencing reads were performed with Trim Galore! (http://www.bioinformatics.babraham.ac.uk/projects/trim_galore/). Co-assembly of the trimmed reads from all samples was performed with MEGAHIT v1.0.2 (Li et al., 2015, 2016) (--kmin-1pass --k-min 27 --k-max 127 --k-step 10).

Contigs >2.5kbp were binned with CONCOCT (Alneberg et al., 2014) while limiting the number of clusters to 400 to avoid ‘fragmentation errors’, within anvi’o v2.3 (Eren et al., 2015) following the metagenomics protocol as previously described. Bins of archaeal origin were manually refined in anvi’o. Completeness and redundancy were estimated with anvi’o single copy gene (SCG) collection, based on Rinke *et al*. (2013), as well as with CheckM v1.0.13 (Parks et al., 2015). In the case of CheckM, analyses were performed by both the standard parameters, with automatic detection of the marker gene set, and by forcing the universal Archaea marker gene collection.

### Phylogenomics and phylogenetics

Prodigal v2.6.3 (-p meta) was used to predict proteins from the archaeal MAGs and 242 archaeal reference genomes (Table S1). Predicted proteins were assigned to archaeal Clusters of Orthologous Groups (arCOGs) using the arNOG database of eggNOG v4.5.1 (Huerta-Cepas et al., 2016) using eggNOG-mapper v0.99.2 (Huerta-Cepas et al., 2017).

Initial phylogenomic reconstruction was based on the 16 ribosomal proteins by Hug *et al*. (2016) with the exception of rpL16 due to frequent misannotation (Laura Hug, pers. comm.) (data not shown) using the Phylosift HMM profiles (Darling et al., 2014). The resulting trees showed relatively poorly resolved basal branches in addition to the absence of rpL22 genes from virtually all DPANN Archaea. We found this to also be the case with other known HMM profile-based databases such as TIGRFAM (Haft et al., 2013). As an example, the rpL14p in DPANN Archaea is often overlooked by the TIGR03673 even when the protein is present as per the more suitable model defined by arCOG04095. Subsequently, we used the more up to date arNOG and increased the number of phylogenetic markers to 44 universal and archaeal ribosomal proteins. Markers were selected from the universal and archaeal ribosomal protein list from Yutin et al. (2012) with the following criteria: i) present in a minimum of 80% of the collection of reference genomes and ii) have a limited level of duplications (standard deviation of copy number in genomes with given arCOG <1.25) (Table S2).

Ribosomal proteins were individually aligned with MAFFT-L-INS-i v7.305b (Yamada et al., 2016) and concatenated. Concatenated alignment was trimmed with BMGE v1.2 (-m BLOSUM30 -g 0.5) (Criscuolo and Gribaldo, 2010) based on Lazar *et al*. (2017).

The concatenated archaeal protein tree was constructed using the posterior mean site frequency (PMSF) model (Wang et al., 2018) in IQ-TREE v1.5.5 (Nguyen et al., 2015), a faster and less resource-intensive approximation to the C10–C60 mixture models (Le et al., 2008), which are also a variant of Phylobayes’ CAT model. Briefly, a base tree was inferred using a simpler substitution model such as LG. This tree was used as the base to infer the final tree under the optimum model suggested by IQ-TREE, LG+C60+F+I+G, with 1000 ultrafast bootstrap replicates (Minh et al., 2013).

Taxonomic identity of the recovered MAGs was also evaluated with GTDB-Tk v 0.3.0 (Chaumeil et al., 2020) using GTDB r89 (Parks et al., 2020). The program was run with both classify_wf and then with de_novo_wf in order to attain a better placement for the MAGs with a limited number of closely related references (Table S3).

#### Phylogenetic placement of new taxa in the DPANN tree

In addition to the rp44 and GTDB-tk analyses, phylogenetic placement of the LFLS DPANN was investigated with additional methods based on a dataset containing all the representative genomes of DPANN from GTDB r89 (151), the 25 LFLS DPANN MAGs, and 10 representatives from Altiarchaeota as outgroup.

Three methods were used for the DPANN-specific phylogenomic analysis: i) concatenated proteins; ii) partitioned analysis; and, iii) multispecies coalescence (MSC) model with ASTRAL. For these, the protein sequences used for tree inference were derived from the 122 SCG used for the archaeal GTDB (ar122) with certain caveats, as 31 proteins were present in <50% of the full dataset (186 genomes). In all cases, each marker was aligned with MAFFT-L-INS-i v7.407 (Yamada et al., 2016).

i) Concatenated protein (dpann-cat): a concatenated MSA was generated as for the rp44 dataset using MAFFT and BMGE. The tree was generated using the PSMF model. Briefly, an initial tree was generated with FastTree v2.1.10 (-lg -gamma) (Price et al., 2010). The best model was searched with IQ-TREE v2.1.2 for all supported mixture models plus iterations of LG with C10–C60 models (Le et al., 2008). The final tree was created with model LG+C60+F+I+G and 1000 ultrafast bootstrap replicates (Hoang et al., 2018).
ii) Partitioned analysis (dpann-part): the MSA derived from the concatenated protein tree was used as input where each partition was defined as per each protein. A total of 91 partitions with a length of 55-749 amino acids were included in the analysis. Substitution models were selected individually per partition. Support values in IQ-TREE were calculated based on 1000 ultrafast bootstrap replicates with nearest neighbour interchange optimisation and --sampling GENESITE.
iii) Supertree with ASTRAL (dpann-astral): sequences for each marker were aligned with MAFFT-L-INS-i and trimmed with BMGE v1.2 with options defined above. Individual ML trees were generated for each marker with IQ-TREE v2.1.2 and the suggested model for each, with 1000 ultrafast bootstrap replicates with nearest neighbour interchange optimisation (Hoang et al., 2018). The resulting ML trees were used as input for ASTRAL-III v5.7.7 (Zhang et al., 2018). This approach was also performed with untrimmed MSA.

In the three scenarios, the markers were derived from the 122 archaeal markers used in the GTDB, with certain caveats, as 31 proteins were present in <50% of the full dataset (186 genomes) and therefore automatically eliminated from the concatenated and partitioned trees.

#### Other phylogenies

Phylogeny of NarG-like proteins was created using all proteins (416 sequences) as reference, from related orthologous groups from the EggNOG database, i.e.: arCOG01497 (19), ENOG4102T1R (5), ENOG4102TDC (5), ENOG4105CRU (351) and ENOG4108JIG (36). Protein sequences were clustered with CD-HIT v4.6 (-c 0.9 -s 0.9 -g 1) (Li and Godzik, 2006) resulting in 276 sequences. Sequences were aligned with MAFFT-L-INS-i v7.305b (Yamada et al., 2016). Alignment was trimmed with BMGE v1.2 (-g 0.9 -h 1 -m BLOSUM30) (Criscuolo and Gribaldo, 2010) and the tree built with IQ-TREE v1.5.5 under the recommended model (LG+R10) and 10,000 ultrafast bootstrap replicates (Nguyen et al., 2015).

Phylogeny of LysJ/ArgD and related proteins was created using all proteins (500 sequences) as reference from related orthologous groups from Swissprot. Protein sequences were clustered with CD-HIT v4.6 (with default parameters) (Li and Godzik, 2006) resulting in 256 representative sequences. Sequences were aligned with MAFFT-L-INS-i v7.305b (Yamada et al., 2016). Alignment was trimmed with BMGE v1.2 (with defaults) (Criscuolo and Gribaldo, 2010) and the tree built with IQ-TREE v1.5.5 under the recommended model (LG+F+I+G4) and 1,000 ultrafast bootstrap replicates (Nguyen et al., 2015).

### Functional annotation

In addition to arCOGs, predicted proteomes were profiled with InterProScan v5.25-64.0 (Jones et al., 2014). InterProScan was run with options --disable-precalc --pathways --goterms -- iprlookup and using the following databases: CDD v3.16 (Marchler-Bauer et al., 2017), Gene3D v4.1.0 (Lees et al., 2014), HAMAP v201701.18 (Pedruzzi et al., 2015), PANTHER v11.1 (Mi et al., 2013), Pfam v31.0 (Finn et al., 2014), PIRSF v3.02 (Wu et al., 2004), PRINTS v42.0 (Attwood et al., 1994), ProDom v2006.1 (Servant et al., 2002), ProSitePatterns and ProSiteProfiles v20.132 (Sigrist et al., 2013), SFLD v2 (Akiva et al., 2014), SMART v7.1 (Letunic et al., 2015), SUPERFAMILY v1.75 (Gough et al., 2001), and TIGRFAM v15.0 (Haft et al., 2013).

Key biogeochemical enzymes were identified based on signature hmm profiles in the InterProScan output or by custom hmm profiles not integrated in databases (Anantharaman et al., 2016; Dombrowski et al., 2017). Custom hmm profiles were searched with HMMER v3.1b2 (-E 1e-20) (Eddy, 2011). Proteins with single HMMER matches below previously established thresholds (Anantharaman et al., 2016; Dombrowski et al., 2017) were searched against Swissprot and checked against the InterProScan results to reduce the incidence of false negatives.

High-heme cytochromes (≥10 binding sites per protein) were predicted using a modified motif_search.py script (https://github.com/alexherns/biotite-scripts) that uses CX(1,4)CH as well as the more canonical CXXCH heme-binding motif.

Carbohydrate-active enzymes were searched locally with hmmscan (HMMER v3.1b2) (Eddy, 2011) and the --domtblout output using a CAZy-based (André et al., 2014) hmm library v5 from dbCAN (Yin et al., 2012). Output was parsed as recommended with the hmmscan-parser.sh script.

Peptidases and proteases were annotated by similarity search of the predicted proteins against the MEROPS database v11.0 (blastp -max_hsps 1 -evalue 1e-4 -seg yes -soft_masking true - lcase_masking -max_target_seqs 1) (Rawlings et al., 2014).

Transporters were annotated by similarity search against the TCDB blastp -max_hsps 1 -evalue 1e-4 -seg yes -soft_masking true -qcov_hsp_perc 50 -lcase_masking -max_target_seqs 1 (database version downloaded on 12 July 2018) (Saier et al., 2014).

The subcellular location of the proteins was predicted with PSORTb v3.0.6 (Yu et al., 2010).

Prediction of tRNA and rRNA genes was performed with tRNAscan-SE 2.0 (Chan et al., 2019) and Barrnap v0.9 (https://github.com/tseemann/barrnap) respectively. Total tRNAs are reported based on the 20 default aminoacyl-tRNAs plus the initiator tRNA (iMet).

### Genome annotation

Metagenome assembled genomes were annotated with Prokka v1.12 (Seemann, 2014) with the --rfam option for the annotation of ncRNA genes. Annotations were imported into PathwayTools v22.0 (Karp et al., 2016) for modelling. Predicted proteins from each MAG were also processed with KofamScan v1.1.0 for KEGG annotation (Kanehisa et al., 2016; Aramaki et al., 2020).

### Multivariate analysis of DPANN genomes

DPANN MAGs from LFLS were compared with extant publicly available genomes based on functional and compositional characteristics. In addition to the genomes in the rp44 dataset, all (165) DPANN published genomes in IMG were downloaded (15 Nov 2019). Genomes from rp44 and IMG were compared with MASH v2.2.2 (Ondov et al., 2016) to remove redundant genomes, resulting in 133 additional DPANN genomes to be added to the dataset (Table S4).

Functional data included in the analysis included COG categories, PSortb predictions, TCDB, MEROPS and CAZy main categories as percentage of the total proteins predicted in each genome. Compositional data included the proportion of each amino acid in the predicted proteome, GC content and coding density. Before analysis, all reference genomes with %C <70% (based on CheckM Archaea) were removed.

Multivariate analysis was performed on zero-centred scaled data via Sparse Principal Component Analysis (sPCA) (Shen and Huang, 2008) as implemented in the MixOmics R package v6.13.21 (Rohart et al., 2017) and limiting the number of explanatory variables per component to five.

### Completeness of DPANN MAGs

Due to the limited completeness values obtained for DPANN genomes – even those in the literature reported to be assembled in single contigs (Table S5), four different Archaeal or prokaryotic SCG collections were examined to assess each of the constituent markers with regard to their suitability for being included as markers for reporting completeness of DPANN MAGs. In addition to the already calculated completeness based on Rinke (default in anvi’o until v5.5 – anvi’o Rinke, 162 SCG) (Eren et al., 2021) and CheckM Archaea (149 SCG), two additional SCG libraries were utilised: anvi’o Archaea76 (76 SCG - current archaeal SCG in anvi’o) (Lee, 2019), and CheckM prokaryote (56 SCG, forced prokaryote/root SCG collections from CheckM).

An initial set of DPANN genomes based on the one used for the multivariate analysis were filtered at 50%C (193 genomes) and 75%C (101 genomes) based on completeness values obtained by CheckM Archaea. All individual markers in all SCG collections were evaluated for their prevalence in each set of genomes. Redundant markers across SCG collections were also compared in order to filter potential universal SCG that might be represented by best/flawed HMM profiles. Initial marker filtering required a prevalence of >=75% across all DPANN genomes, and a prevalence within classic DPANN lineages of >50% to account for the limited number of highly complete genomes.

### Pangenomic analysis

Pangenomic analysis of the found ‘*Ca.* Methanoperedens spp.’ genomes, as well as the four nearly complete reference ‘*Ca.* Methanoperedens spp.’ was performed with anvi’o v5.4.0 (Eren et al., 2015) following the standard pangenomics workflow (http://merenlab.org/2016/11/08/pangenomics-v2). The reference genomes included in all comparisons and pangenome analysis were: ‘*Ca.* Methanoperedens nitroreducens’ ANME-2D (type genome, JMIY00000000.1) (Haroon et al., 2013), ‘*Ca.* Methanoperedens ferrireducens’ (PQAS00000000.1) (Cai et al., 2018), ‘*Ca.* Methanoperedens batavicum’ comb. nov. (previously referred to as ‘*Ca.* M. nitroreducens’ BLZ1 or MPEBLZ) (LKCM00000000.1) (Arshad et al., 2015), and ‘*Ca.* Methanoperedens vercellense’ comb. nov. (previously ‘*Ca.* M. nitroreducens’ Vercelli, GCA_900196725.1) (Vaksmaa et al., 2017). Protein-coding genes were clustered with an MCL inflation value of 6 (van Dongen and Abreu-Goodger, 2012).

Average Nucleotide Identity (ANI) and Average Amino acid Identity (AAI) were calculated with pyani v0.2.7 (Pritchard et al., 2015) and CompareM v0.0.23 (https://github.com/dparks1134/CompareM) respectively.

### Proposed taxa

Details on etymology and nomenclature can be found in the Taxonomic Appendix. A summary of the proposed taxa can be found in Table 1.

**Table 1.**
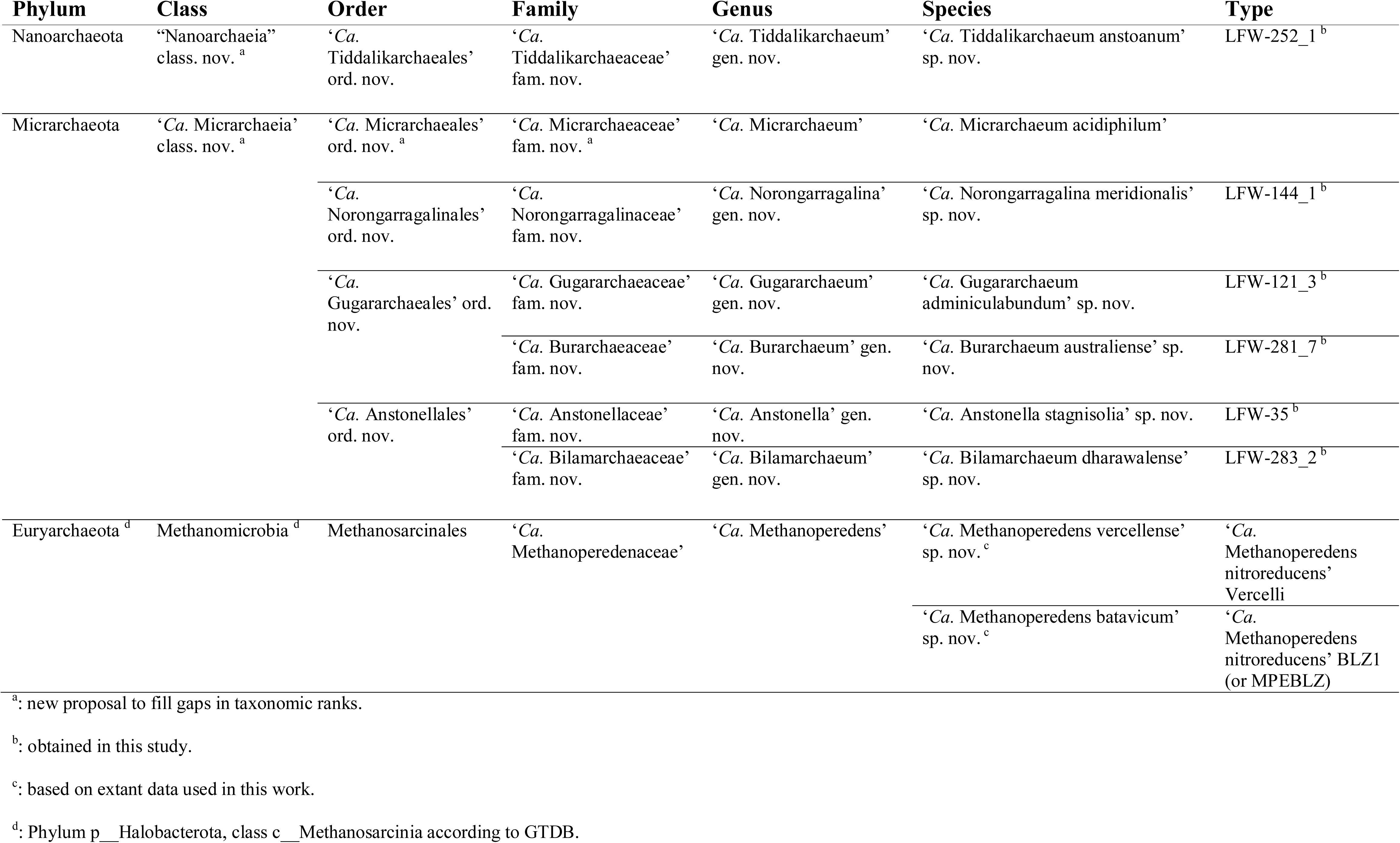
Summary of new taxa proposed. Formal descriptions of the taxa in this table are included in the Taxonomic Appendix with additional details on the proposed type materials in Table S6.

Minimal quality criteria for denominating new candidate species was based on Chuvochina et al. (2019). Relaxed parameters were allowed for selected genomes when phylogenetic/taxonomic novelty was deemed of special relevance. Under those conditions no new names were proposed even under taxonomic novelty (e.g. LFWA-IV) for MAGs with completeness <90%, <18 tRNA, and/or 16S rRNA gene sequence <1000 bp. Additional details about the proposed species can be found in Table S6.

### Data availability

Annotated assemblies are available at ENA under project PRJEB21808 and sample identifiers ERS2655284–ERS2655320. See details in Table S7. The output of the different analyses, including the reference genomes, are available at Zenodo (doi:10.5281/zenodo.3365725).

## Results

### Binning results and archaeal community

In order to better understand the archaeal contributions in LFLS groundwaters and their influence upon site biogeochemistry, raw sequencing reads from ENA project PRJEB14718 (Vázquez-Campos et al., 2017) were reanalysed using genome-based metagenomic methodologies. Briefly, samples were collected in triplicate over a period of 47 days at four time-points (0, 4, 21 and 47 days) after an intense rainfall event that filled the trench. Thorough chemical and radiochemical analyses were conducted on the samples in order to fully characterise the geochemistry (Vázquez-Campos et al., 2017). Over this time period, the redox conditions transitioned from slightly oxic to increasingly reducing/anoxic, reflected in both the chemical analyses and the community profile, with a clear increase in obligate anaerobes on days 21 and 47 (Vázquez-Campos et al., 2017).

The *de novo* co-assembly of sequence reads from the LFLS trench subsurface water samples (Vázquez-Campos et al., 2017) generated 187,416 contigs with a total length of 1.32 Gbp (based on 2.5 kbp cutoff). Binning with CONCOCT generated 290 initial bins. A total of 21 bins with clear archaeal identity or with ambiguous identity (similar completeness scores for bacterial and archaeal SCG profiles) were further refined with anvi’o, producing 37 curated archaeal MAGs with completeness ≥50% (%C from here on; 22 of ≥90%C) and redundancy ≤10% (%R from here on) (Table 2 and Table S8 for detailed completeness assessment).

**Table 2.**
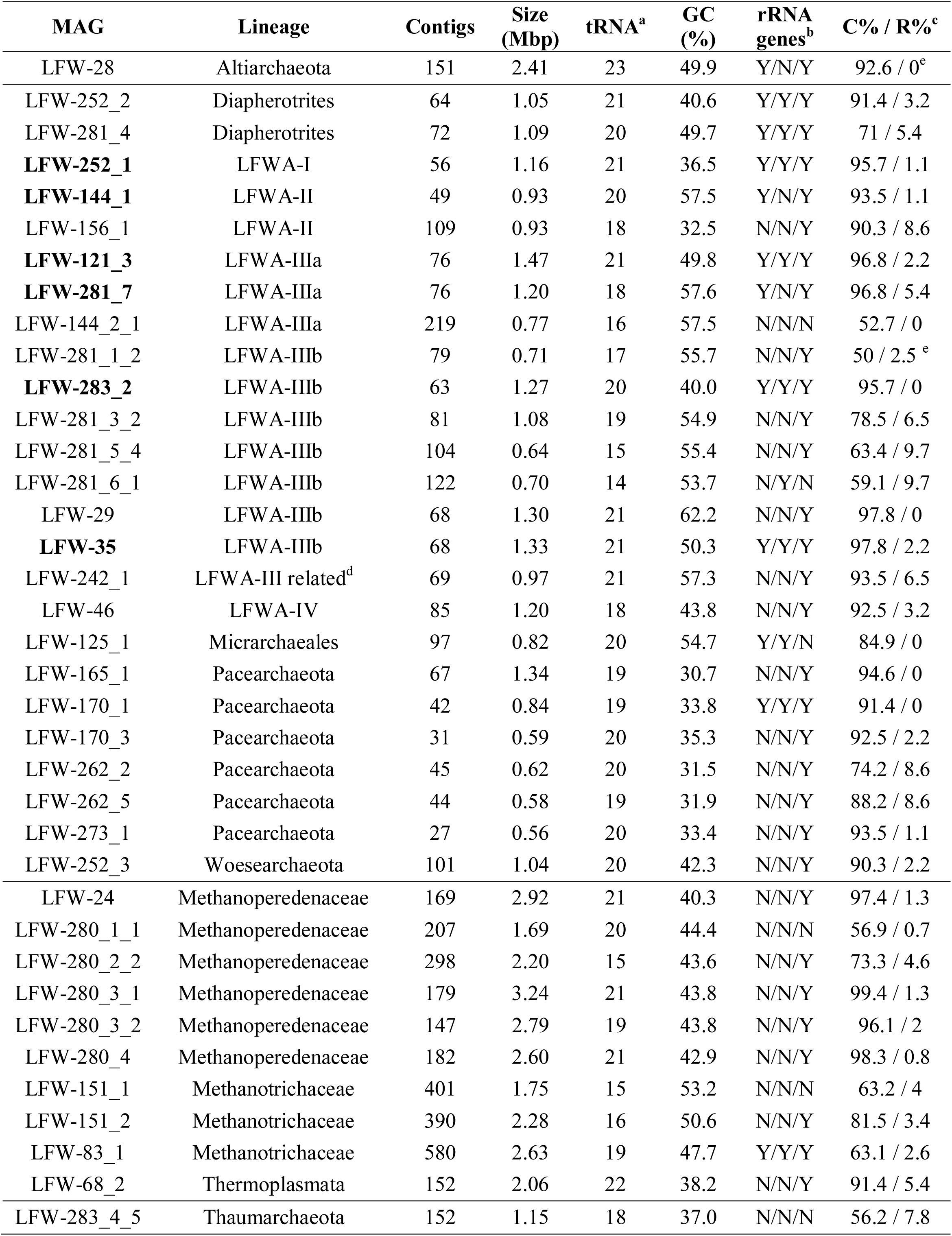
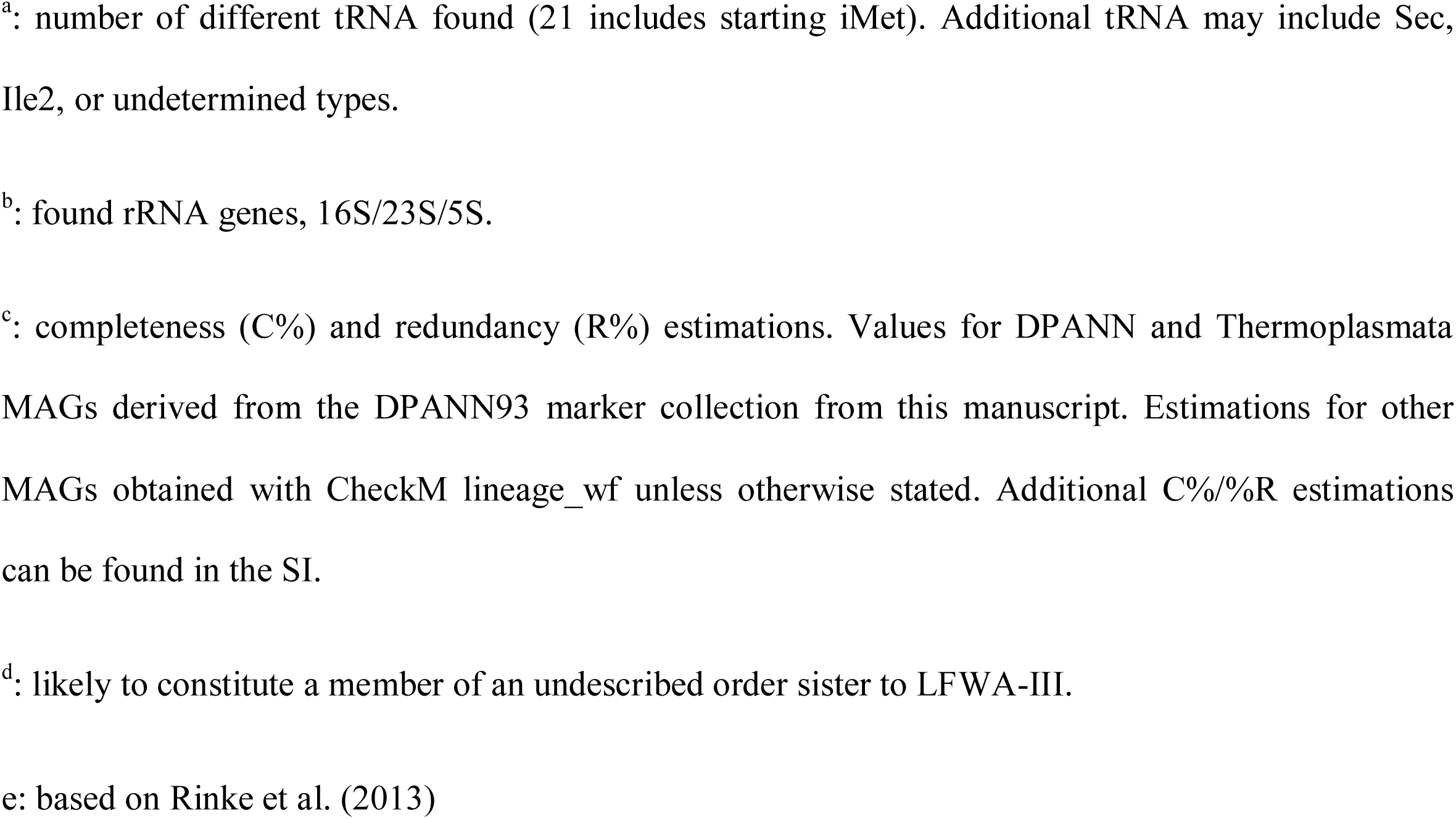
General characteristics of the genomes reconstructed in this study. MAGs in bold indicate named candidate species (see Table 1). Additional details can be found in Table S7. Expanded completeness analysis are available in Table S8.

| MAG | Lineage | Contigs | Size (Mbp) | tRNA<sup>a</sup> | GC (%) | rRNA genes<sup>b</sup> | C% / R%<sup>c</sup> |
| --- | --- | --- | --- | --- | --- | --- | --- |
| LFW-28 | Altiarchaeota | 151 | 2.41 | 23 | 49.9 | Y/N/Y | 92.6 / 0<sup>e</sup> |
| LFW-252_2 | Diapherotrites | 64 | 1.05 | 21 | 40.6 | Y/Y/Y | 91.4 / 3.2 |
| LFW-281_4 | Diapherotrites | 72 | 1.09 | 20 | 49.7 | Y/Y/Y | 71 / 5.4 |
| **LFW-252_1** | LFWA-I | 56 | 1.16 | 21 | 36.5 | Y/Y/Y | 95.7 / 1.1 |
| **LFW-144_1** | LFWA-II | 49 | 0.93 | 20 | 57.5 | Y/N/Y | 93.5 / 1.1 |
| LFW-156_1 | LFWA-II | 109 | 0.93 | 18 | 32.5 | N/N/Y | 90.3 / 8.6 |
| **LFW-121_3** | LFWA-IIIa | 76 | 1.47 | 21 | 49.8 | Y/Y/Y | 96.8 / 2.2 |
| **LFW-281_7** | LFWA-IIIa | 76 | 1.20 | 18 | 57.6 | Y/N/Y | 96.8 / 5.4 |
| LFW-144_2_1 | LFWA-IIIa | 219 | 0.77 | 16 | 57.5 | N/N/N | 52.7 / 0 |
| LFW-281_1_2 | LFWA-IIIb | 79 | 0.71 | 17 | 55.7 | N/N/Y | 50 / 2.5<sup>e</sup> |
| **LFW-283_2** | LFWA-IIIb | 63 | 1.27 | 20 | 40.0 | Y/Y/Y | 95.7 / 0 |
| LFW-281_3_2 | LFWA-IIIb | 81 | 1.08 | 19 | 54.9 | N/N/Y | 78.5 / 6.5 |
| LFW-281_5_4 | LFWA-IIIb | 104 | 0.64 | 15 | 55.4 | N/N/Y | 63.4 / 9.7 |
| LFW-281_6_1 | LFWA-IIIb | 122 | 0.70 | 14 | 53.7 | N/Y/N | 59.1 / 9.7 |
| LFW-29 | LFWA-IIIb | 68 | 1.30 | 21 | 62.2 | N/N/Y | 97.8 / 0 |
| **LFW-35** | LFWA-IIIb | 68 | 1.33 | 21 | 50.3 | Y/Y/Y | 97.8 / 2.2 |
| LFW-242_1 | LFWA-III related<sup>d</sup> | 69 | 0.97 | 21 | 57.3 | N/N/Y | 93.5 / 6.5 |
| LFW-46 | LFWA-IV | 85 | 1.20 | 18 | 43.8 | N/N/Y | 92.5 / 3.2 |
| LFW-125_1 | Micrarchaeales | 97 | 0.82 | 20 | 54.7 | Y/Y/N | 84.9 / 0 |
| LFW-165_1 | Pacearchaeota | 67 | 1.34 | 19 | 30.7 | N/N/Y | 94.6 / 0 |
| LFW-170_1 | Pacearchaeota | 42 | 0.84 | 19 | 33.8 | Y/Y/Y | 91.4 / 0 |
| LFW-170_3 | Pacearchaeota | 31 | 0.59 | 20 | 35.3 | N/N/Y | 92.5 / 2.2 |
| LFW-262_2 | Pacearchaeota | 45 | 0.62 | 20 | 31.5 | N/N/Y | 74.2 / 8.6 |
| LFW-262_5 | Pacearchaeota | 44 | 0.58 | 19 | 31.9 | N/N/Y | 88.2 / 8.6 |
| LFW-273_1 | Pacearchaeota | 27 | 0.56 | 20 | 33.4 | N/N/Y | 93.5 / 1.1 |
| LFW-252_3 | Woesearchaeota | 101 | 1.04 | 20 | 42.3 | N/N/Y | 90.3 / 2.2 |
| LFW-24 | Methanoperedenaceae | 169 | 2.92 | 21 | 40.3 | N/N/Y | 97.4 / 1.3 |
| LFW-280_1_1 | Methanoperedenaceae | 207 | 1.69 | 20 | 44.4 | N/N/N | 56.9 / 0.7 |
| LFW-280_2_2 | Methanoperedenaceae | 298 | 2.20 | 15 | 43.6 | N/N/Y | 73.3 / 4.6 |
| LFW-280_3_1 | Methanoperedenaceae | 179 | 3.24 | 21 | 43.8 | N/N/Y | 99.4 / 1.3 |
| LFW-280_3_2 | Methanoperedenaceae | 147 | 2.79 | 19 | 43.8 | N/N/Y | 96.1 / 2 |
| LFW-280_4 | Methanoperedenaceae | 182 | 2.60 | 21 | 42.9 | N/N/Y | 98.3 / 0.8 |
| LFW-151_1 | Methanotrichaceae | 401 | 1.75 | 15 | 53.2 | N/N/N | 63.2 / 4 |
| LFW-151_2 | Methanotrichaceae | 390 | 2.28 | 16 | 50.6 | N/N/Y | 81.5 / 3.4 |
| LFW-83_1 | Methanotrichaceae | 580 | 2.63 | 19 | 47.7 | Y/Y/Y | 63.1 / 2.6 |
| LFW-68_2 | Thermoplasmata | 152 | 2.06 | 22 | 38.2 | N/N/Y | 91.4 / 5.4 |
| LFW-283_4_5 | Thaumarchaeota | 152 | 1.15 | 18 | 37.0 | N/N/N | 56.2 / 7.8 |

<sup>a</sup>: number of different tRNA found (21 includes starting iMet). Additional tRNA may include Sec, Ile2, or undetermined types.
<sup>b</sup>: found rRNA genes, 16S/23S/5S.
<sup>c</sup>: completeness (C%) and redundancy (R%) estimations. Values for DPANN and Thermoplasmata MAGs derived from the DPANN93 marker collection from this manuscript. Estimations for other MAGs obtained with CheckM lineage\_wf unless otherwise stated. Additional C%/R estimations can be found in the SI.
<sup>d</sup>: likely to constitute a member of an undescribed order sister to LFWA-III.
<sup>e</sup>: based on Rinke et al. (2013)

Phylogeny based on the 44 concatenated ribosomal proteins (rp44, Figure 1) showed that most metagenome assembled genomes (MAGs) from LFLS belonged to diverse DPANN lineages and *Methanomicrobia* (*Methanoperedenaceae* and *Methanotrichaceae*) (Table 2). Phylogenomic analysis based on the rp44 (Figure 1) showed a general tree topology largely consistent with current studies, e.g. *Asgardarchaeota* as sister lineage to TACK and *Altiarchaeota* as sister to DPANN (Zaremba-Niedzwiedzka et al., 2017; Dombrowski et al., 2019). Most high level branches showed UF bootstrap support values >90%, with the exception of the very basal Euryarchaeota, which is known to be difficult to resolve (Adam et al., 2017). The position of the MAGs in the archaeal phylogeny suggests four new lineages (Figure 1) within DPANN, denoted LFWA-I to -IV. These lineages correspond to representatives of high-level taxa (order or above) that do not have either a current proposed name or have not been explored in detail in the literature. As a means of honouring the Australian Aboriginal community, and very especially, the traditional owners of the land where LFLS is located, many of the nomenclatural novelties proposed are based on terms from Aboriginal languages related to the site (mainly Dharawal), see Taxonomic Appendix. A summary of the nomenclatural proposals derived from this work are summarised in Table 1.

**Figure 1.**
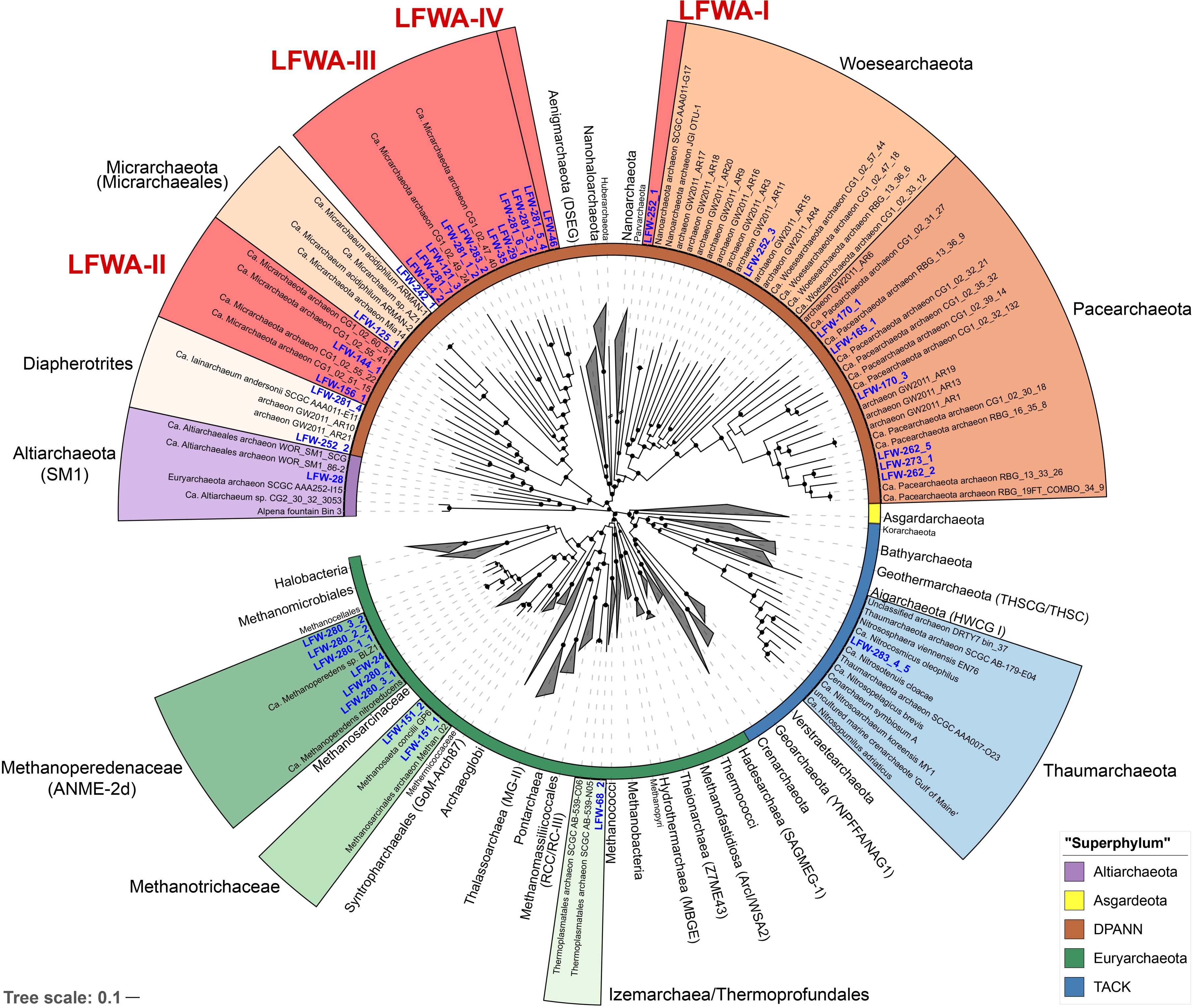
Phylogenomic analysis of the archaeal MAGs found at LFLS. Concatenated protein tree constructed with 44 universal and Archaea-specific ribosomal proteins from 230 reference genomes and 36 original MAGs from this work. Tree was constructed with IQ-TREE under the PMSF+LG+C60+F+I+G. Lineages with representatives at LFLS are shown coloured and uncollapsed, with labels in blue. Inner ring indicates the major archaeal lineages. Shortened branches are shown at 50% of their length. Black circles indicate ultrafast bootstrap support values >90%.

During the drafting of this manuscript, a genome-based taxonomy was developed for Bacteria and Archaea (GTDB) (Parks et al., 2018). For all proposed lineages, we have included the possible correspondence with the GTDB taxonomy (r89) wherever necessary using the GTDB nomenclature with prefixes, e.g. “p ” for phylum, to avoid confusion with more “typically” defined lineages with the same names.

### The archaeal community in detail

Based on bin coverage information, the archaeal community was dominated by DPANN, esp. *Pacearchaeota* and LFWA-III (see below), and *Euryarchaeota,* esp. *Methanoperedenaceae*, with maximum relative abundance values of 79.7% at day 4, and 42.8% at day 47, respectively (Table S9). Our previous analysis indicated that DPANN constituted a maximum of 55.8% of the archaeal community (Vázquez-Campos et al., 2017). This discrepancy could be an artefact derived from the different copy numbers of rRNA gene clusters, often limited to one in DPANN and TACK, compared to other Archaea with more typically sized genomes containing multiple copies of the operon.

Based on the functional annotation of key biogeochemical proteins (Figure 2 and Table S10), the archaeal MAGs recovered from LFLS do not appear to play a major role in the sulfur cycle (lacking dissimilatory pathways, Table S10). Regarding the nitrogen cycle, the presence of nitrogenases in some of the *Methanoperedens* MAGs (Figure 2 and Table S10) suggest that they may play a larger role in the fixation of N_2_ than in the dissimilatory reduction of nitrate, particularly given the negligible nitrate concentrations measured (Vázquez-Campos et al., 2017). Another Archaea likely to be heavily involved in the nitrogen cycle, but through the degradation of proteins, is the *Thermoplasmata* LFW-68_2, based on the unusually high number of proteases encoded in its genome (Figure 2, Figure S1 and Table S11; see SI for details).

**Figure 2.**
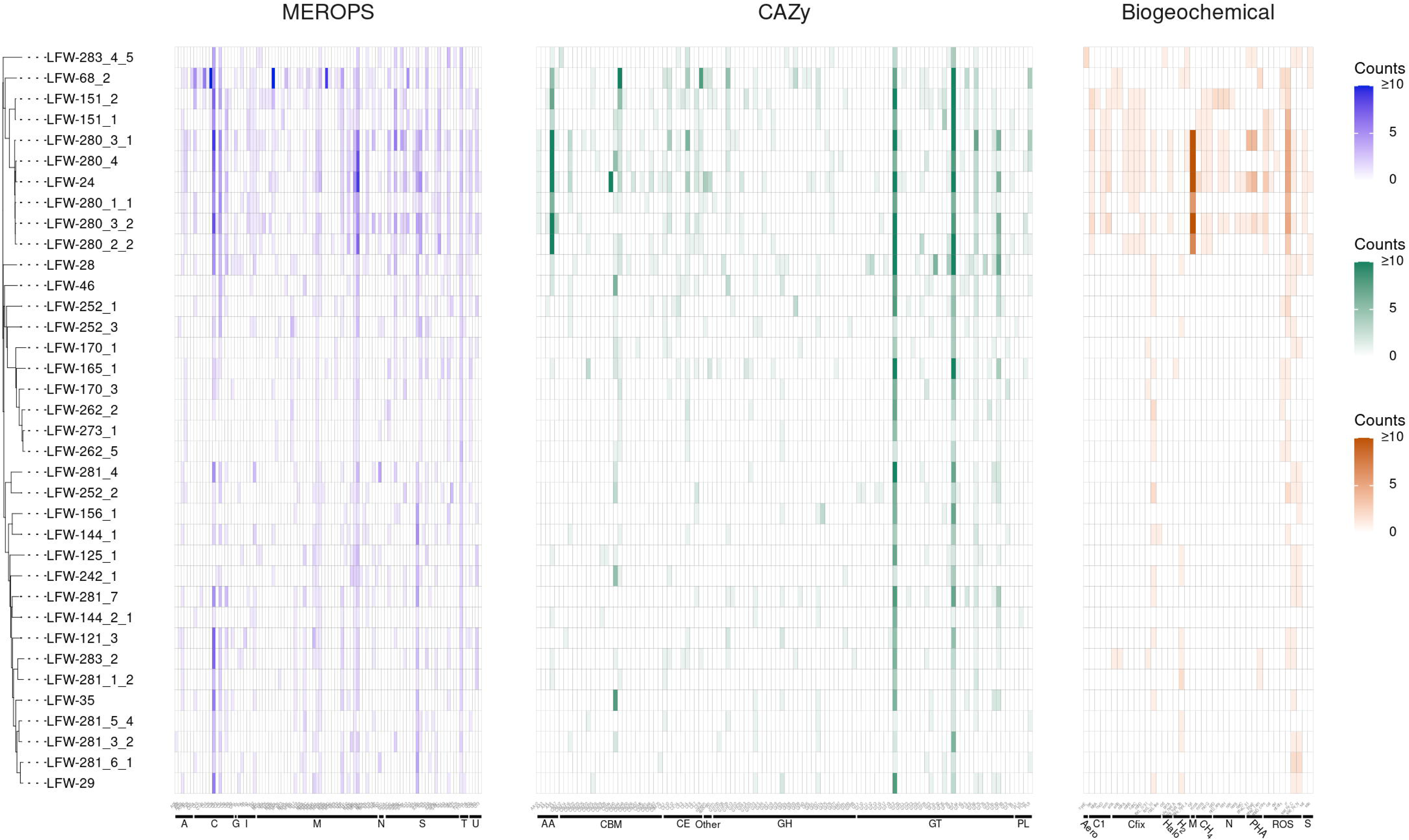
Abundance of MEROPS subfamilies, CAZy families, and biogeochemically relevant proteins found in the MAGs from LFLS. MAGs are ordered according to the rp44 tree (Figure 1). Counts of each protein are capped at 10 for easier visualisation. Details about MEROPS, CAZy, and proteins of biogeochemical relevance, including exact counts and classes with predicted extracellular proteins can be found in Tables S11, S13 and S10, respectively. Plot created with ggtree v3.1.0 (Yu, 2020) and ggtreeExtra v1.0.4 (Xu et al., 2021).

Due to their special interest and relevance, the content below is focused on the new DPANN and the *Methanoperedens* spp. from LFLS. Aspects related to other MAGs and other general observations regarding the whole archaeal community are contained within the Supplementary Materials.

### DPANN in Little Forest Legacy Site

#### LFWA-I: candidate order *Tiddalikarchaeales*

The LFWA-I lineage is a sister lineage to *Parvarchaeota* (ARMAN-4 and -5) (Baker et al., 2006; Rinke et al., 2013) in rp44 and contains a unique MAG, LFW-252_1 (‘*Ca.* Tiddalikarchaeum anstoanum’) (Figure 1, Figure S5). LFWA-I corresponds to the order and family o CG07-land f CG07-land within the c *Nanoarchaeia* as per GTDB. As with many other DPANN from the PANN branch (such as cluster 2 in Dombrowski et al (2020)) %C estimates with existing Archaea SCG collections are severely underestimated with barely 79.0%C reported by CheckM (best). The %C combined with the assembly size sets a complete genome size >1.5 Mbp, well above the expected genome size for a Nanoarchaeota, and similar to that for many PANN MAGs (Table S8). A dedicated SCG collection was built from extant HMM profiles by removing SCG’s normally absent from all DPANN and/or from specific sub-lineages. This first attempt of a tailored SCG library for assessing the completeness of DPANN genomes (Table S12), provides more sensible results not only for LFW-252_1 (95.7%C/1.1%R) but also for DPANN assemblies in general, including some well-studied references (Table S5).

The LFW-252_1 lacks the biosynthetic capacity for almost all amino acids, and for the *de novo* enzymes biosynthesising purine and pyrimidine, despite having a normal enzymatic representation for interconversion of purine nucleotides, and interconversion of pyrimidine nucleotides (Figure S2). With the additional lack of a proper pentose biosynthesis pathway and only containing the last enzyme required for the biosynthesis of 5-phospho-α-D-ribose 1-diphosphate (PRPP), LFW-252_1 is very likely to depend on a host, maybe aerobic or microaerophilic, although this is only based on the relative abundance patterns. Reactive oxygen species detoxification is mediated by superoxide reductase (SOR, neelaredoxin/desulfoferredoxin; TIGR00332) and rubredoxin. LFW-252_1 is a subsurface decomposer based on the CAZymes associated to the degradation of lignocellulosic material, including carbohydrate esterases (CE1, CE12 and CE14), β-mannase (GH113), and even one vanillyl-alcohol oxidase (AA4) (Figure 2, Table S13). Its genome also harbours three different sialidases (GH33), at least two extracellular, making it one of the few non-pathogenic prokaryotes containing this enzyme (Buelow et al., 2016). In addition to the sialic acids as sources of C and N, LFW-252_1 also contains two extracellular serine peptidases (S08A, subtilisin-like), indicating a possible active role in the degradation of proteins in the environment (Table S11). Energy production is based on the ferredoxin-mediated oxidative decarboxylation of pyruvate (PorABC), 2-oxoglutarate (KorAB), and, likely, other oxoacids (OforAB), i.e. fermentation of carbohydrates and amino acids.

#### LFWA-II: candidate order *Norongarragalinales*

The lineage LFWA-II is represented in LFLS by MAGs LFW-144_1 and LFW-156_1, and it is consistently placed as a sister clade to LFWA-III+*Micrarchaeales* (see taxonomic novelties; Figure 1, Figure S2). Both MAGs have a similar size but different completeness estimates and GC content: 93.5%C/1.1%R and 57.5% GC for LFW-144_1, and 90.3%C/8.6%R and 32.5% GC for LFW-156_1. Each of the LFWA-II MAGs belong to different genera based on AAI, and to different families within o UBA8480 (c *Micrarchaeia*) based on the GTDB. ‘*Ca.* Norongarragalina meridionalis’ LFW-144_1, is the most complete genome within this order and co-generic with the only representative on the placeholder family f 0-14-0-20-59-11.

Functionally, LFW-144_1 is an auxotroph for most amino acids, displaying little capacity for amino acid interconversion (Figure S3). Conversely, it carries a complete pentose-phosphate pathway and a capacity for *de novo* biosynthesis of purine and pyrimidine nucleotides. While it lacks extracellular CAZymes (Figure 2), it has an extracellular metallopeptidase M23B (lysostaphin-like). M23 peptidases are often used to break bacterial cell walls as either a defence mechanism (bacteriocin) or for nutrition (e.g. predation or parasitism). Energy production is derived from the fermentation of pyruvate (PorABCD).

Despite the similar assembly size (<3 kbp difference) and similar %C compared to LFW-144_1, LFW-156_1 presents an even more limited genome, lacking complete *de novo* nucleotide biosynthesis and pentose phosphate pathways. Conversely, it has three different extracellular glycoside hydrolases (GH74, GH57, GH23) and one protease (S08A). It ferments D-lactate (D-lactate dehydrogenase, LDH) instead of pyruvate.

#### LFWA-III is two distinct but similar *Micrarchaeota* lineages: candidate orders *Gugararchaeales* and *Anstonellales*

LFWA-III is the closest branch to *Micrarchaeales*, constituting a match with GTDB’s placeholding order o UBA10214 within c *Micrarchaeia* according to the GTDB-tk output. The lineage LFWA-III was the most diverse of all Archaea encountered at LFLS with 10 MAGs belonging to three different families and 10 different genera based on AAI values and phylogeny (Tables S14 and S15).

Phylogenetic analysis and RED values suggest though that LFWA-III is actually two similar but distinct orders (Figure 3). The RED for the two possible orders (LFWA-IIIa, 0.526; and LFWA-IIIb, 0.525) are closer to the median value to the RED estimates for order (0.43–0.57, median 0.51) than the LFWA-III’s RED (0.47). While LFWA-IIIb seems to be topologically consistent across trees, LFWA-IIIa is paraphyletic in rp44 and dpann-astral. In addition, both the GTDB-tk derived trees and our DPANN trees show LFWA-IIIb to be equivalent to a pre-existing order in GTDB, o UBA10214 (Figure 3).

**Figure 3.**
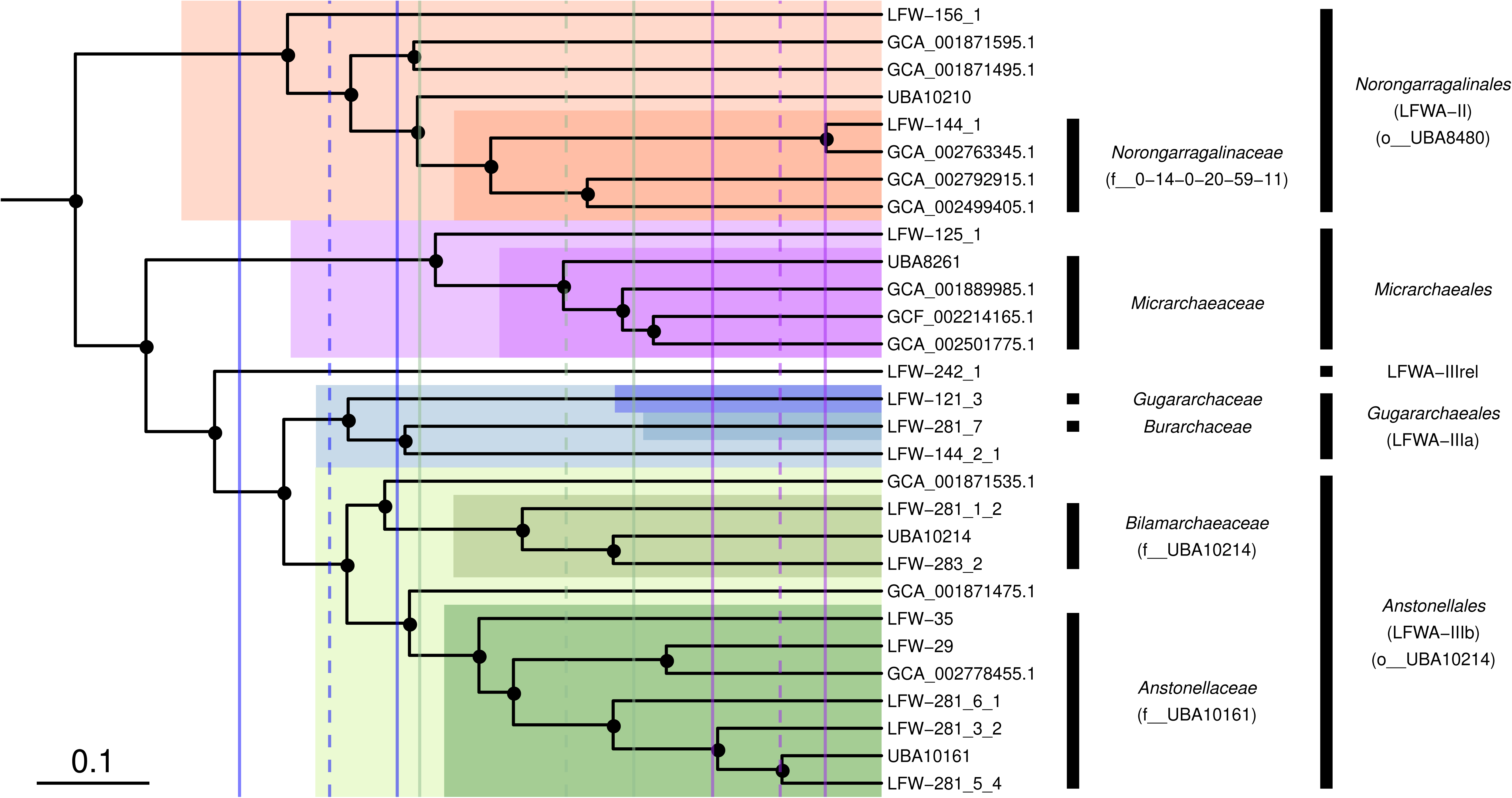
Phylogenomic placement of the *Micrarchaeota* orders and families with respect to the type family *Micrarchaeaceae*. Subtree extracted from the concatenated DPANN phylogenomic tree highlighting families and orders. Filled circles indicate UFBoot support ≥ 90% in the dpann-concat tree. Scale bar indicates RED. Solid vertical lines indicate minimum and maximum RED values for order (blue), family (green), and genus (purple). Dashed lines indicate median RED values.

LFWA-IIIa, or *Gugararchaeales*, includes three MAGs (Table 2) belonging to three different families, but only two of high quality: LFW-121_3, ‘*Ca.* Gugararchaeum adminiculabundum’ LFW-121_3 (96.8%C, 2.2%R, 49.8% GC), and LFW-281_7, ‘*Ca.* Burarchaeum australiense’ LFW-281_7 (96.8%C, 5.4%R, 57.6% GC). Both MAGs constitute the types of their respective families, *Gugararchaeaceae* and *Burarchaeaceae*. On the other hand, LFWA-IIIb or *Anstonellales*, is the most diverse order in LFLS waters, with 7 MAGs from two different families, but only two of high quality: LFW-35, ‘*Ca.* Anstonella stagnisolia’ (97.8%C, 2.2%R, 62.2% GC), and LFW-283_2, ‘*Ca.* Bilamarchaeum dharawalense’ (95.7%C, 0%R, 40.0% GC). *Anstonellaceae* would be equivalent to f UBA10161, while *Bilamarchaeaceae* would be equivalent to f UBA10214. In terms of abundance, LFWA-III, in general, constituted >25% of the archaeal community at any individual sampling day with ‘*Ca.* Anstonella stagnisolia’ LFW-35 being the most abundant Archaea at day 0 and the second most abundant during the remaining days (Table S9).

#### Comparative functional analysis

In addition to their high abundance and diversity, LFWA-III is also an interesting lineage at a functional level, constituting one of the few DPANN with broad anabolic capabilities including the *de novo* biosynthesis of the amino acids lysine, arginine, and cysteine as well as purine and pyrimidine nucleotides (Figure 4). The metabolic capabilities of the top five LFWA-III genomes (the four named species plus LFW-29) were compared with five of the better characterised and/or complete DPANN genomes (Figure 4): ‘*Ca.* Iainarchaeum andersonii’ (Youssef et al., 2015) and AR10 (Castelle et al., 2015) (*Diapherotrites*), ‘*Ca.* Micrarchaeum acidiphilum’ ARMAN-2 (Baker et al., 2006) and ‘*Ca.* Mancarchaeum acidiphilum’ Mia14 (Golyshina et al., 2017) (*Micrarchaeota*), and “*Nanoarchaeum equitans*” Kin4-M (Huber et al., 2002; Waters et al., 2003) (“*Nanoarchaeota*”). In terms of core metabolism, the representative genomes of LFWA-III share more similarities with *Diapherotrites* than with *Micrarchaeota*, e.g. *de novo* biosynthesis pathways for nucleotides, de novo biosynthesis of nicotinate/nicotinamide or aromatic amino acids (Figure 4). However, LFWA-III also contains other unique features: pathways for the biosynthesis of L-arginine, L-lysine, L-cysteine, and L-histidine (only in AR10 in the reference genomes). Both LFWA-IIIa MAGs display most of the genes encoding the pathway for the biosynthesis of branched amino acids. Conversely, in the case of LFWA-IIIb, and likely also with ‘*Ca.* Iainarchaeum andersonii’, would likely rely on the interconversion of L-valine and L-leucine. LFW-121_3 in particular presents unique features that either distinguish it from other LFWA-III genomes and/or the reference genomes. Both LFW-121_3 and ‘*Ca.* Iainarchaeum andersonii’ are the only MAGs able to convert L-aspartate to L-threonine based upon current evidence.

**Figure 4.**
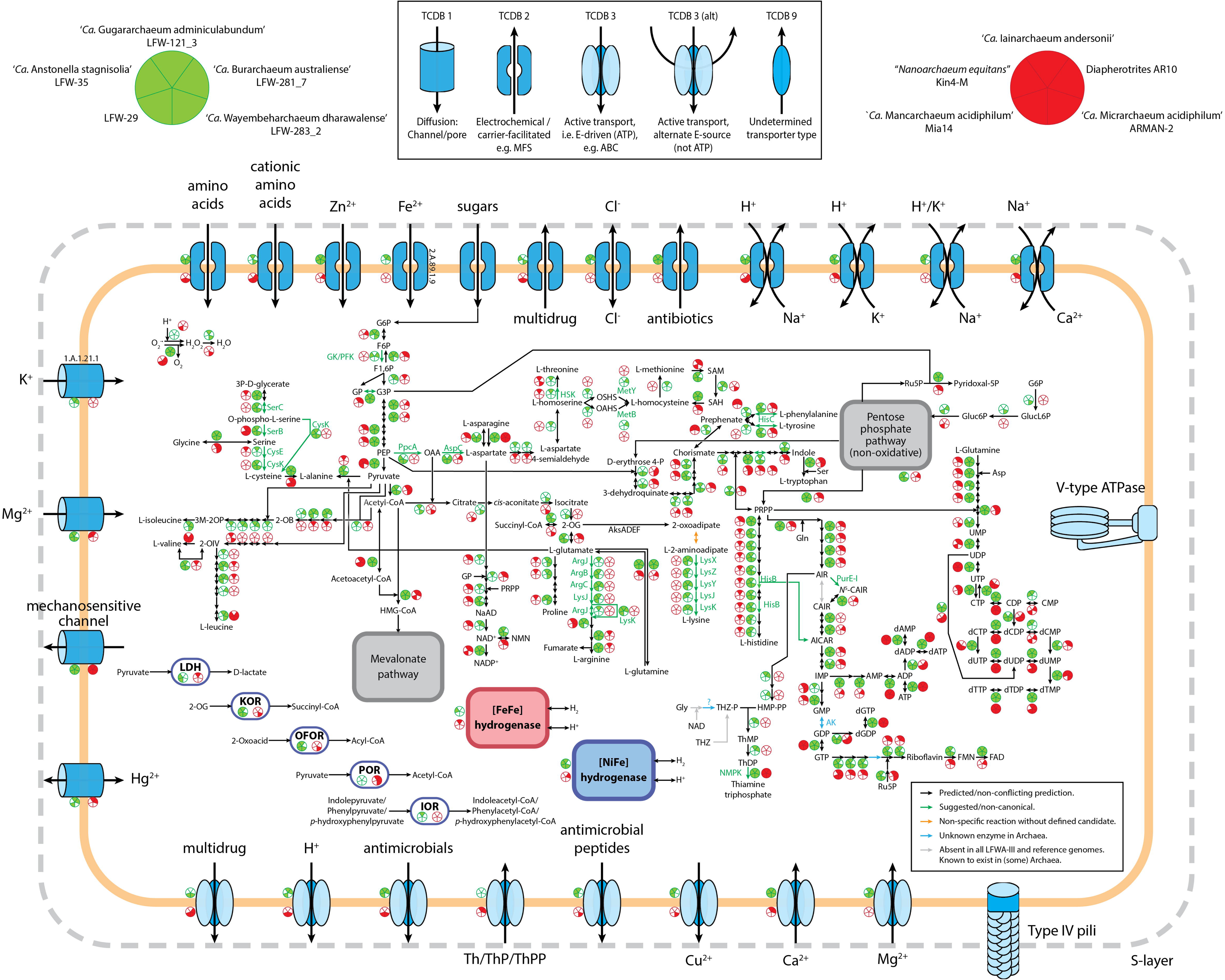
Comparative visualisation of the metabolism of LFWA-III representative MAGs from this study (green circles) and selected reference DPANN genomes (red circled). Each sector in the indicative circles indicates the presence or absence of a key protein in the represented genome. Extracellular enzymes are not shown for simplicity. Abbreviations: 2-OG, 2-oxoglutarate; 2-OIV, 2-oxoisovalerate; AICAR, 5-amino-1-(5-phospho-D-ribosyl)imidazole-4-carboxamide; AIR, 5-amino-1-(5-phospho-β-D-ribosyl)imidazole; ARP, 5-amino-6-ribitylamino-2,4(1H,3H)-pyrimidinedione; CAIR, 5-amino-1-(5-phospho-D-ribosyl)imidazole-4-carboxylate; F1,6P, β-D-Fructose 1,6-bisphosphate; F6P, β-D-Fructose-6P; FAD, Flavin adenine dinucleotide; FMN, Flavin mononucleotide; G3P, Glyceraldehyde-3P; G6P, Glucose-6P; Gluc6P: D-gluconate 6-phosphate; GlucL6P: D-glucono-1,5-lactone 6-phosphate; GP, Glycerone-P; HMP-PP, 4-amino-5-hydroxymethyl-2-methylpyrimidine diphosphate; IOR: indolepyruvate:ferredoxin oxidoreductase; KOR: 2-oxoglutarate:ferredoxin oxidoreductase; LDH: D-lactate dehydrogenase (NAD^+^); N5-CAIR, 5-(carboxyamino)imidazole ribonucleotide; NAD^+^, β-nicotinamide adenine dinucleotide; NADP^+^, β-nicotinamide adenine dinucleotide phosphate; NMN, β-nicotinamide D-ribonucleotide; OAA, Oxaloacetate; OAHS, O-acetyl-homoserine; OFOR: 2-oxoacid:ferredoxin oxidoreductase; OSHS, O-succinyl-homoserine; PEP, Phosphoenolpyruvate; POR: pyruvate:ferredoxin oxidoreductase; PRPP, 5-phospho-α-D-ribose 1-diphosphate; Ru5P, D-ribulose 5-phosphate; SAH, S-adenosyl-L-homocysteine; SAM, S-adenosyl-L-methionine; ThDP, Thiamine diphosphate; ThMP, Thiamine monophosphate; THZ, 4-methyl-5-(β-hydroxyethyl)thiazolium; THZ-P, 4-methyl-5-(β-hydroxyethyl)thiazolium phosphate.

Another interesting difference between *Diapherotrites* and LFWA-IIIa relates to thiamine acquisition (Figure 4). Both *Diapherotrites* genomes have transporters for thiamine and/or its phosphorylated forms but lack any enzyme known in the *de novo* pathway for its biosynthesis. Instead, LFWA-IIIa is likely to be able to produce thiamine *de novo* provided a source for the thiazole branch of the thiamine pathway is available (see SI for further discussion). None of the LFWA-IIIb MAGs have the biosynthetic pathway nor specific transporters for thiamine.

Like other DPANN, all LFWA-III are fermentative organisms. However, there are differences in the substrates used. Leaving aside LFW-281_7, for which we couldn’t detect any of the proteins mentioned next, all other LFWA-III and ARMAN-2 can ferment 2-oxoglutarate (KorAB), and other 2-oxoacids (OforAB). Pyruvate fermentation was detected in both *Diapherotrites* and ARMAN-2 but not in any LFWA-III. *Anstonellaceae* MAGs (LFW-35, LFW-29) were the only ones with LDH, a differential characteristic from the other LFWA-III, but shared with ‘*Ca.* Iainarchaeum andersonii’. LFW-121_3 was the only one with a predicted capacity for the fermentation of aromatic amino acids amongst the 10 genomes compared (IorAB).

Regarding transporters (Figure 4), the high variability across individual MAGs makes generalisations difficult. Nonetheless, certain types of transporters show a distinctive distribution. Chloride channel proteins (TCDB 2.A.49.6.1), TCDB-2 antibiotic exporters (e.g. MarC family), H^+^,K^+^/Na^+^ antiporters and Mg^2+^ P-type ATPase (MgtA/MgtB, Mg^2+^ import) appear unique to LFWA-III. Two transporters appear to be specific for LFWA-IIIa: the potassium channel protein (TCDB 1.A.1.21.1) and the uncharacterised TCDB 1.B.78.1.4 now annotated as a VIT family protein (TCDB 2.A.89.1.9), often involved in iron homeostasis. Ca^2+^/Na^+^ and H^+^/K^+^ antiporters, and Ca^2+^ P-type ATPase are common to both LFWA-III and *Diapherotrites*.

#### LFWA-IV: an *incertae sedis* DPANN lineage

The MAG LFW-46 is the sole representative of the LFWA-IV lineage (Figure 1). While presenting good %C/%R levels for a functional description (92.5%C, 3.2%R, 43.8% GC), it lacks all rRNA genes aside from a partial 5S (thus no name is formally proposed, Figure S3). All the DPANN trees place LFW-46 as sister to EX4484-52 (member of the unnamed homonym phyla) and, together, a sister to *Nanohaloarchaeota*.

The LFW-46 genome is a typical example of a reduced, host-dependent DPANN (Figure S4) with very limited amino acid interconversion pathways, lacking both the *de novo* nucleotide biosynthesis as well as the pentose phosphate pathways. However, LFW-46 differentiates itself by the relatively rich repertoire of transporters related to resistance, especially in comparison with the repertoire of other DPANN: multidrug, tetracycline, quinolones, arsenite, Cu^+^ and Cu^2+^, and mono-/di-valent organocations (e.g. quaternary ammonium compounds). The extracellular enzymes of LFW-46 include one protease (S08A), one glycoside hydrolase (GH57), and one polysaccharide lyase (PL9). LFW-46 ferments pyruvate for energy (PorABCD).

#### Multivariate comparison of DPANN genomes

Sparse Principal Component Analyses (sPCA) of the functional (e.g. COG, and MEROPS annotations) and/or compositional (amino acid composition of predicted proteins, and GC%) profiles of selected Archaea and DPANN genomes (only ≥75%C based on CheckM) were performed in order to reveal the overall distinctiveness of LFWA lineages. In the all-Archaea dataset with compositional+functional data, LFWA-III clustered closer to other DPANN genomes (Figure S5) yet positioned near to what could be referred to as a ‘transition zone’ between DPANN and normal-sized archaeal genomes (e.g. Euryarchaeota and TACK). Compositional and functional analysis provided an insight into the uniqueness of LFWA-III: displaying high GC% (usually ∼50% or greater) and an overall above-average number of COG-annotated proteins.

Functional sPCA of the 140 DPANN genomes enabled differentiation of the metabolically rich (e.g. LFWA-II, LFWA-III, Diapherotrites and related) and acidophilic lineages from the other DPANN genomes (Figures 5A and 5B). This differentiation of LFWA-III was mainly shown in PC1 (Figure 5A; COG categories L, O, U, W) and PC3 (not shown; COG E, F, H and MEROPS C).

**Figure 5.**
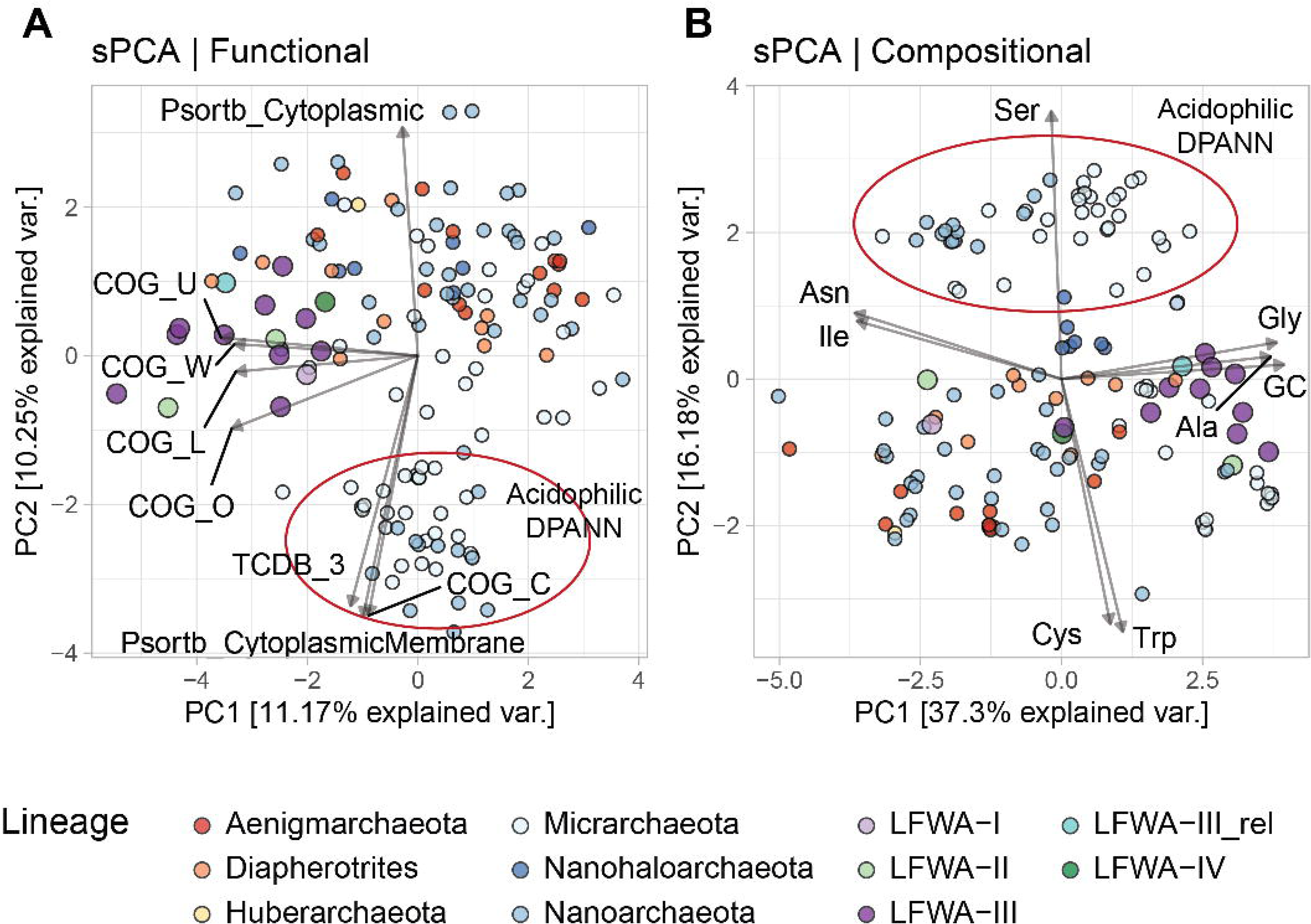
sPCA analysis of the functional (A) and compositional (B) features in DPANN genomes. Only explanatory variables correlated at absolute value of ≥ 0.75 with the principal components are shown. Variables are scaled (4x) for easier visualisation. MAGs from LFLS lineages are displayed with larger circles.

The high positive loadings in the compositional dataset along PC1 for Gly, Ala, and GC, and negative loadings for Ile and Asn may be due to the high GC content of most LFWA-III genomes (Table 2, Figure 5B), considering that those amino acids are encoded by GC-rich and AT-rich codons respectively (Grosjean and Westhof, 2016). Very high positive values along PC2 separated the acidophilic DPANN related to the acidophilic *Micrarchaeota* and *Parvarchaeota* from the other DPANN.

### Diverse and abundant ‘*Ca.* Methanoperedens spp.’

The occurrence of three methanogenic MAGs from the *Methanotrichaceae* family (previously indicated to be the illegitimate *Methanosaetaceae* (Vázquez-Campos et al., 2017)) were principally and, unsurprisingly, detected during the highly anoxic phase (117–147 mV Standard Hydrogen Electrode). More interestingly though, was the recovery of MAGs belonging to the genus ‘*Ca.* Methanoperedens’. A total of six MAGs were assembled in this study, four ≥95%C (all <10%R). Each of these MAGs belong to different species based on ANI and AAI scores (Figure 6, Tables S16 and S17), constituting the most diverse set of *Methanoperedens* spp. genomes reconstructed to date in a single study and site, and all different to previously sequenced MAGs.

**Figure 6.**
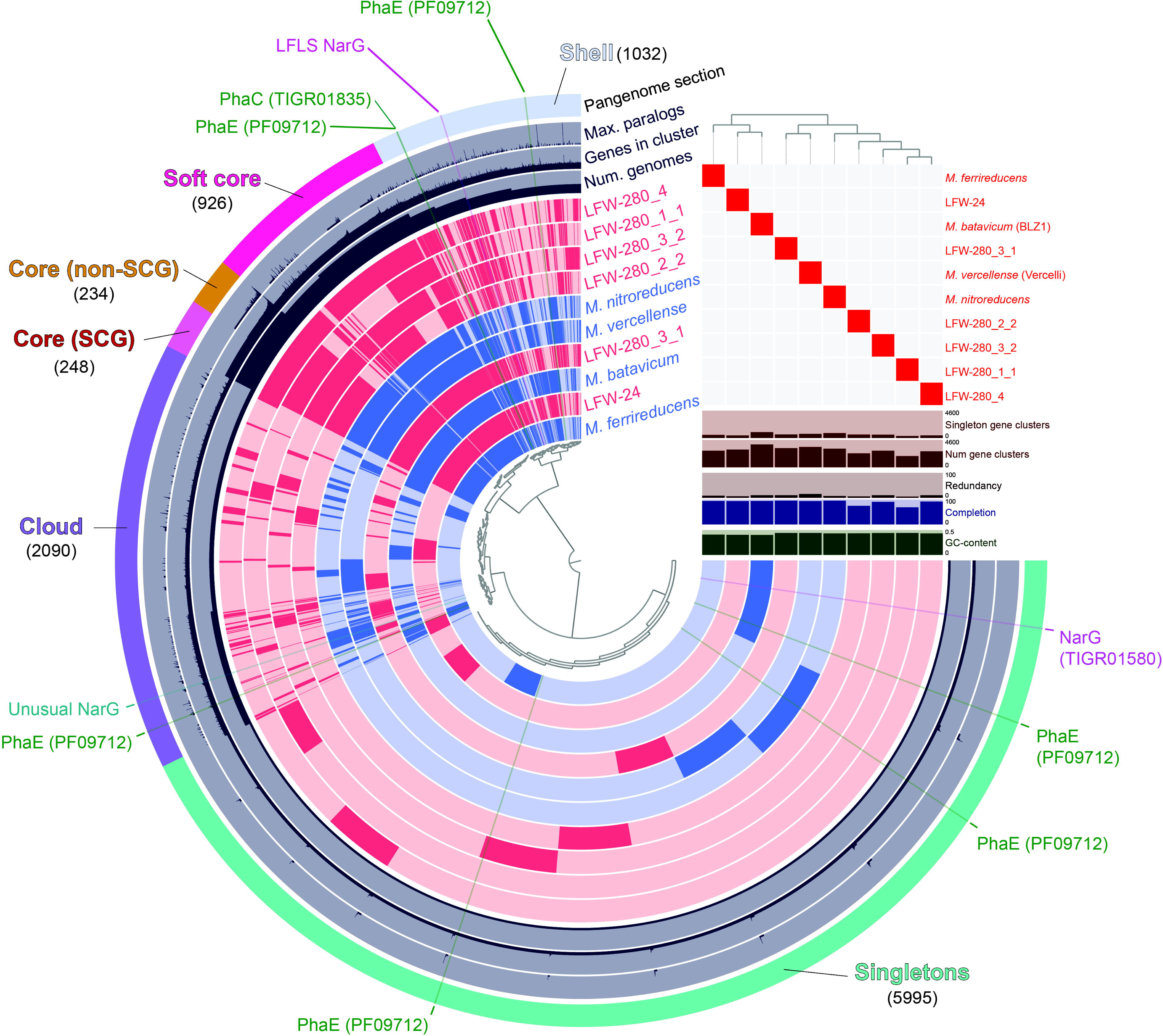
Pangenomic analysis of genus *Methanoperedens*. Pangenome was generated with 6 *Methanoperedens* spp. MAGs from LFLS (pink) and 4 reference genomes (blue), accounting for a total of 31,974 protein-coding genes grouped in 10,525 gene clusters. Gene clusters generated with an MCL inflation value of 6.0.

The rich collection of *Methanoperedens* spp. recovered, in combination with selected published genomes, allows for the universal investigation of common aspects across the whole genus. A total of four reference genomes are included in all comparisons and pangenome analysis: ‘*Ca.* Methanoperedens nitroreducens’ ANME-2D (Haroon et al., 2013), ‘*Ca.* Methanoperedens ferrireducens’ (Cai et al., 2018), ‘*Ca.* Methanoperedens batavicum’ comb. nov. (previously referred to as ‘*Ca.* M. nitroreducens’ BLZ1 or MPEBLZ) (Arshad et al., 2015), and ‘*Ca.* Methanoperedens vercellense’ comb. nov. (previously ‘*Ca.* M. nitroreducens’ Vercelli) (Vaksmaa et al., 2017) (see Taxonomic appendix and SI).

#### Multiple acquisition of respiratory molybdopterin oxidoreductases

Most known *Methanoperedens* spp. encode some molybdopterin oxidoreductase enzyme, which are usually referred to as NarG. All the *Methanoperedens* spp. found at LFLS share a putative molybdopterin oxidoreductase not present in other members of the genus (Figure 7). Notably, this is different from any canonical or putative NarG (as is often suggested for other *Methanoperedens* spp.) with no match to HMM profiles characteristic of these proteins but, instead, belonging to arCOG01497/TIGR03479 (DMSO reductase family type II enzyme, molybdopterin subunit).

**Figure 7.**
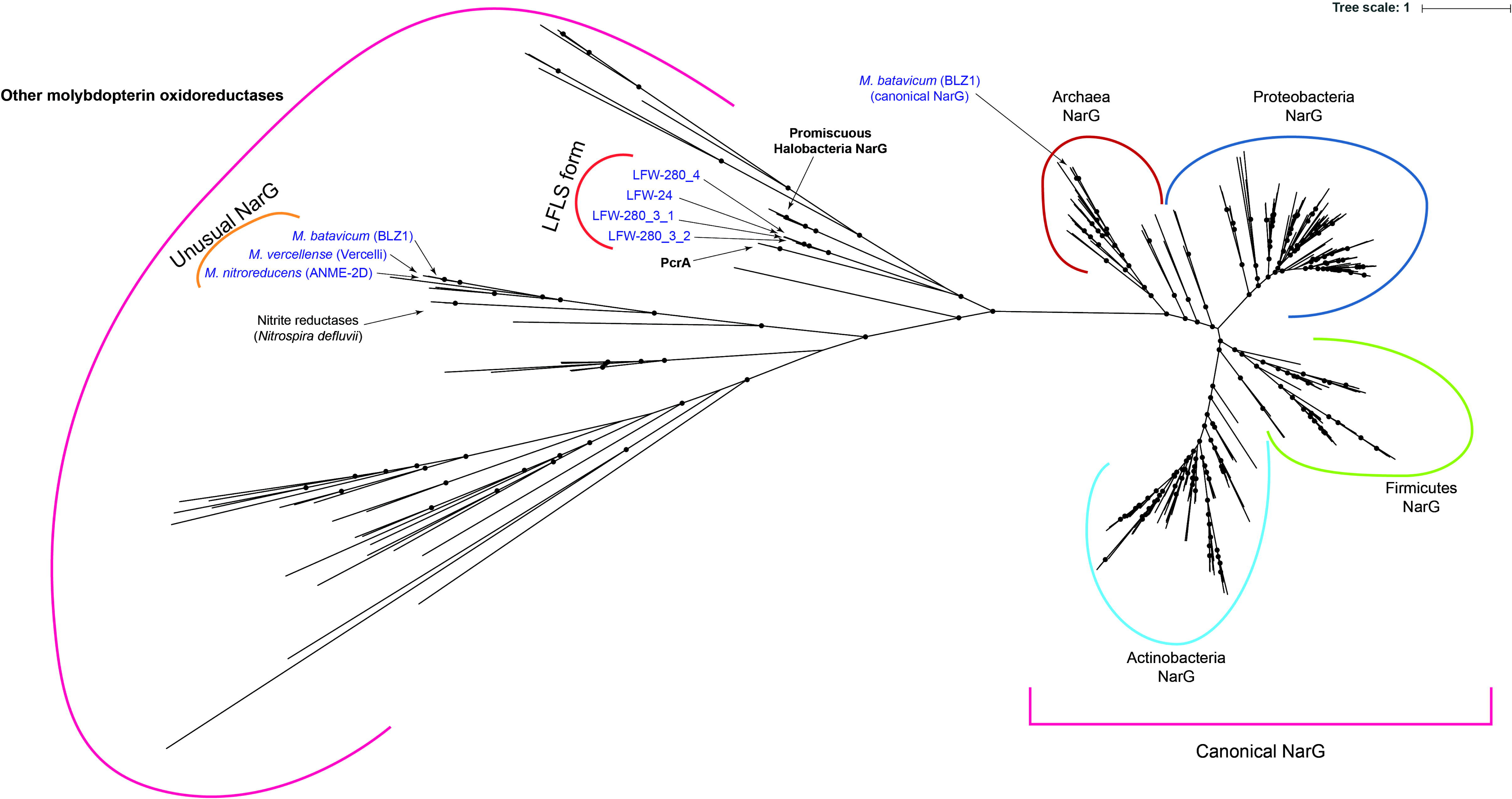
Phylogenetic analysis of molybdopterin oxidoreductase proteins. Tree was generated with 268 reference sequences and 8 query sequences from ‘*Ca.* Methanoperedens’ spp. NarG(-like) proteins. Tree was built under model LG+R10 model and 10,000 ultrafast bootstrap iterations. Branches with ultrafast bootstrap support values <50% are collapsed. Black circles indicate a branch support ≥90%.

Phylogenetic analysis (Figure 7) places the molybdopterin oxidoreductase form of LFLS *Methanoperedens* spp. far from molybdopterin oxidoreductases from the reference *Methanoperedens* spp.. However, it is located closer to non-canonical NarG sequences from *Haloferax mediterranei* (I3R9M9) (Lledó et al., 2004) and other *Halobacteria*, as well as to the perchlorate reductase (PcrA) of *Dechloromonas aromatica* and *Azospira oryzae* (previously known as *Dechlorosoma suillum* strain PS) (He et al., 2019).

#### High heme cytochromes are common in *Methanoperedens* spp. genomes

Metal respiration is usually associated with cytochromes with a high number of heme-binding sites (≥10) or high heme cytochromes (HHC from here on). The screening of the *Methanoperedens* spp. MAGs from LFLS revealed multiple HHC in all genomes, with a minimum of six copies for LFW-280_1_1 (68.5%C, Figure 2, Table S10). All reference *Methanoperedens* spp. genomes revealed similar abundances when considering their completeness, i.e., ∼10 HHC per genome. While most gene clusters containing HHC are present in the cloud genome or as singletons, there is one highly conserved gene cluster in the core genome (GC_00000135) of *Methanoperedens* containing 13 heme-binding sites. Although metal reduction has only been confirmed for *M. ferrireducens* and *Methanoperedens batavicum* BLZ1 (Ettwig et al., 2016; Cai et al., 2018), our data suggests metal reduction might be a universal characteristic of *Methanoperedens* spp.

#### Biosynthesis of polyhydroxyalkanoates

Polyhydroxyalkanoates (PHA) are carbon-rich, energetic polymers that result from the polymerisation of hydroxyalkanoates, e.g. 3-hydroxybutyrate, produced by numerous microorganisms under unfavourable conditions, particularly in association with unbalanced nutrient levels (Reddy et al., 2003; Martínez-Gutiérrez et al., 2018). Genes involved in the biosynthesis of polyhydroxyalkanoates (PHA) were detected in three of the most complete *Methanoperedens* spp. genomes, i.e. LFW-24, LFW-280_3_1 and LFW-280_3_2, as well as in the *Thaumarchaeota* LFW-283_4_5 (Figure 2, Table S10).

In *Methanoperedens* spp., the genes associated with the biosynthesis of PHA are usually present as an operon (Figure S6A). Although not explicitly reported in their respective manuscripts, PhaC (PHA synthase type III heterodimeric, TIGR01836) and PhaE (PHA synthase subunit E, PF09712) were predicted in all the *Methanoperedens* reference genomes included in this study (Haroon et al., 2013; Arshad et al., 2015; Vaksmaa et al., 2017; Cai et al., 2018). However, while all PhaC proteins form a gene cluster in the pangenome analysis, PhaE does not (Figure 6). This could be related to their function: PhaC is a catalytic protein while PhaE ‘simply’ regulates PhaC (Koller, 2019), so any sequence modifications in PhaC might have a more critical effect on the biosynthesis of PHA. Both proteins, PhaC and PhaE, have also been predicted in three of the most complete new genomes in the present study (LFW-24, LFW-280_3_1 and LFWA-280_3_2). In most instances, PHA biosynthesis genes have been found in putative operons (Figure S8A).

## Discussion

In this study we describe 22 new DPANN Archaea candidate taxa (species to order), as well as proposing four supraspecific taxa to accommodate extant and newly proposed lineages according to formal taxonomy (family to class), along with the proposal of two ‘*Ca.* Methanoperedens spp.’ based on extant data.

The archaeal community at LFLS, showed a clear dominance of DPANN and ‘*Ca.* Methanoperedens spp.’, with the data sets’ uniqueness highlighted by the unmatched MAGs (at species level) with any extant reference genomes. Amongst the DPANN MAGs recovered, a number of them constitute the first detailed description of lineages with placeholder names from large scale projects (Parks et al., 2017, 2020; Rinke et al., 2020).

### Self-sustaining DPANN – possibly more common than previously thought?

The DPANN archaea *sensu stricto* (i.e. not including *Altiarchaeota*) are often described as organisms with extremely reduced metabolic capacities and generally, with few exceptions, dependent on a host for the acquisition of essential metabolites such as vitamins, amino acids or even reducing equivalents (Dombrowski et al., 2019). During the metabolic reconstruction of individual archaeal genomes, it was observed that select DPANN from the LFWA-III lineage, exemplified by LFW-121_3 (‘*Ca.* Gugararchaeum adminiculabundum’), possess near-complete pathways for the synthesis of amino acids, purine and pyrimidine nucleotides, riboflavin, and thiamine (Figure 4). These predictions, derived from either Pathway Tools (via Prokka annotations) or KEGG, were unusual given that when a DPANN genome lacks the biosynthetic capacity for an amino acid, all enzymes for that pathway are typically missing. However, several of the pathways in the *Gugararchaeales* and *Anstonellales* MAGs had a limited number of gaps (Figure 4). These alleged ‘gaps’, thoroughly discussed in the SI with focus on LFW-121_3, were often related to unusual enzymes poorly annotated in databases, or steps known to be possibly performed by bifunctional or promiscuous enzymes. Manual curation with predicted functions from other databases combined with literature searches was able to fill some of those gaps or, at least, provide reasonable candidate proteins that may perform those functions.

The discovery of additional DPANN taxa with a “rich” metabolism, at least compared with host-dependant DPANN (e.g. “*Nanoarchaeum*”, *Nanopusillus*, Mia14), highlights the metabolic diversity and widespread existence of free-living DPANN *Archaea*, at least within the *Diapherotrites*/*Micrarchaeota* lineage – DM branch or cluster 1 (Dombrowski et al., 2020) – as opposed to the PANN branch, suggesting a more generalised genome streamlining process/loss of function in the latter. Prior studies have remarked upon the generalised “limited metabolic capacities” in most DPANN (Dombrowski et al., 2019). However, many DPANN datasets are heavily enriched with lineages that are likely to be host-dependent, e.g. *Pacearchaeota* or *Woesearchaeota* (suggested to be orders in the GTDB), or derived from acidic environments, generally restricted to ARMAN lineages, esp. *Micrarchaeaceae*. This study expands the phylum *Micrarchaeota* with 3 orders, 5 families, 5 genera and 5 species. (Figure 3).

Several studies have reported that ultra-small *Bacteria* and *Archaea* can be lost using standard 0.22 µm filters (Luef et al., 2015; Chen et al., 2018). However, this may only affect individual cells, while host-DPANN cell associations would be retained and overrepresented compared to free-living DPANN, which are not necessarily larger than the host-associated organisms. The high abundance of the putative free-living LFWA-III in the metagenomic samples from LFLS could be related to the elevated concentrations of labile colloidal Fe that precipitates at the redox interface following the infiltration of oxic-rainwaters. Acting in a similar manner as a flocculant in water treatment, iron (oxyhydr)oxide microaggregates have immense capacity to trap organic matter and microbial cells (Tang et al., 2016). This process has been used in protocols to recover and concentrate viruses from environmental samples without the need of ultrafiltration (John et al., 2011). As such, the colloidal Fe would also have improved the recovery of free-living DPANNs and any other ultra-small microorganisms that, by default would have been otherwise lost during the filtration with the 0.22 µm filters used during sampling (Vázquez-Campos et al., 2017).

### ‘*Ca.* Methanoperedens’: molybdopterin oxidoreductases, metal reduction and PHA biosynthesis

*Methanoperedenaceae* archaea have been reported (or at least suggested) to be able to utilise a wide range of inorganic electron acceptors for AOM, including nitrate (Haroon et al., 2013), nitrite (Arshad et al., 2015), Fe(III) (Ettwig et al., 2016; Cai et al., 2018), Mn(IV) (Ettwig et al., 2016) and Cr(VI) (Lu et al., 2016). The type species of the genus *Methanoperedens*, ‘*Ca.* Methanoperedens nitroreducens’, receives its name from its ability to use nitrate as an electron acceptor (Haroon et al., 2013). However, the alpha subunit of this candidate respiratory nitrate reductase (NarG_Mn_) is a molybdopterin oxidoreductase different from any canonical or putative NarG, with no match with TIGRFAM, CDD or Pfam profiles characteristic of these proteins. This NarG_Mn_ does not belong to the same orthologous group alongside other typical archaeal NarG (arCOG01497), but to ENOG4102T1R, which includes the well-characterised “dimethyl sulfoxide reductase subunit A” (DmsA) of several *Halobacteria*. The NarG_Mn_ has a high similarity (>80%) with proteins from both *Methanoperedens batavicum* BLZ1 (NarG1_MB_) and *Methanoperedens vercellense* (Vercelli, NarG_MV_) and relates to the nitrite reductases of *Nitrospira defluvii* (Figure 7). While *Methanoperedens vercellense* and *M. nitroreducens* have only this unusual NarG, *Methanoperedens batavicum* harbours an additional, unrelated, canonical NarG (NarG2_MB_) based on the detection by the TIGRFAM profile (TIGR01580, arCOG01497).

Several of the NarG related to NarG_LFLS_ (from *Halobacteria*) have been confirmed to have certain promiscuity (Figure 7) and are known to be able to use (per)chlorate as a substrate *in vivo* (Yoshimatsu et al., 2000; Oren et al., 2014). Although it is close to impossible to predict the actual substrates of the NarG_LFLS_ (or PcrA_LFLS_), the phylogenetic analysis is, at least, suggestive of the potential utilisation of (per)chlorate. The oxidation of methane linked to the bioreduction of perchlorate has been proposed a number of times in the past (Luo et al., 2015; Xie et al., 2018; Wu et al., 2019). Recent works (Xie et al., 2018; Wu et al., 2019) have suggested the possible involvement of ANME archaea in (per)chlorate reduction based on chemical and amplicon data analyses, an assumption considered more likely given that nitrate was rarely measured in noticeable concentrations in the LFLS sampling trench (Vázquez-Campos et al., 2017). Indeed, nitrate concentrations were below the detection limit (<0.01 μM) before any *Methanoperedens* became relatively abundant. With LFLS historical disposal records indicating that quantities of perchlorate/perchloric acid were deposited in the vicinity of the sample location, it collectively suggests that *Methanoperedens* spp. utilisation of nitrate is unlikely at the site. However, as no chlorite dismutase gene was found in any of the MAGs, the possibility that they may utilise perchlorate is also not well supported. Given the high iron concentrations, it is reasonable to assume that the AOM at LFLS uses Fe(III) as the main electron acceptor.

The collective evidence indicates that nitrate reductase(-like) enzymes are rather widespread in *Methanoperedens* spp., with the exception of ‘*Ca.* M. ferrireducens’ from which no nitrate reductase candidate could be detected, aside from an orphan NapA-like protein (cd02754). Phylogenetic analysis of the NarG and similar proteins (Figure 7) suggests that these proteins might have been horizontally acquired at least three different times during the evolution of *Methanoperedens*. Congruently, the pangenomic analysis showed three different NarG variants in their respective gene clusters (Figure 6 and Figure 7).

#### High-heme cytochromes

Analysis of historical disposal records revealed that over 760 (intact and partially corroded) steel drums were deposited within the legacy trenches at LFLS, including in close proximity of the collected samples (Payne, 2012). The unabated redox oscillations which have occurred in the trenches over the last 60+ years and the resultant impact upon the steel drums, have likely contributed to the elevated concentrations of soluble iron observed (∼0.5–1 mM) (Vázquez-Campos et al., 2017).

Utilisation of metals (e.g. Fe, Mn, Cr, U) as electron acceptors requires the presence of multi-heme cytochromes. Although multi-heme cytochromes are not intrinsically and exclusively employed for heavy metal reduction (e.g. decaheme DmsE for DMSO reduction), HHC are characteristic of microorganisms performing biological reduction of metals. In our case, HHC were found in all new and reference *Methanoperedens*, including a gene cluster from the core genome. This may well indicate that oxidised metal species were the ancestral electron acceptor to all *Methanoperedens*, rather than nitrate or other non-metallic oxyanion moieties.

Previous studies indicate that *M. batavicum* and *M. ferrireducens* can both use the Fe(III) (oxyhydr)oxide ferrihydrite as electron acceptor for AOM (Ettwig et al., 2016; Cai et al., 2018). Reports showing the involvement of more crystalline forms of iron oxides for the AOM are scarce with the possibility that these organisms may drive a cryptic sulfate reduction cycle rather than a more direct Fe(III)-dependent AOM (Sivan et al., 2014). Based on the evidence to hand, it is not clear whether *Methanoperedenaceae* archaea or other ANME could be responsible for these observations.

Tantalising questions remain as to the role of *Methanoperedens* spp. with regard to the ultimate fate of iron within the legacy trenches; an important consideration given the central role that iron plays in the mobilisation/retardation of key contaminants plutonium and americium (Ikeda-Ohno et al., 2014; Vázquez-Campos et al., 2017).

#### PHA biosynthesis

Variable nutrient levels can result in the microbial production of polyhydroxyalkanoates (PHA), which are carbon-rich, energetic polymers resulting from the polymerisation of hydroxyalkanoates (Reddy et al., 2003; Martínez-Gutiérrez et al., 2018). The biosynthesis of PHA is a widespread characteristic in many groups of aerobic *Bacteria* (Reddy et al., 2003; Martínez-Gutiérrez et al., 2018). However, very few strict anaerobes are able to synthesise PHA, being mostly limited to syntrophic bacteria (Spang et al., 2012). In *Archaea*, the biosynthesis of PHA is known to occur, particularly in many *Nitrososphaerales* (*Thaumarchaeota*) (Spang et al., 2012) and *Euryarchaeota*, where it has been traditionally limited to *Halobacteria* (Koller, 2019).

Key genes, as well as full operons, have been detected in several *Methanoperedens* genomes, with recent experimental evidence identifying the production of PHA by *Methanoperedens nitroreducens* (Cai et al., 2019). This suggests that the biosynthesis of PHA could be a widespread feature in *Methanoperedens* spp. or even in all *Methanoperedenaceae*, indicating a possible, generalised, role in the accumulation of excess carbon at times when the carbon source (methane) is much more abundant than other nutrients and/or trace elements (Karthikeyan et al., 2015).

It has been estimated that anaerobic methane oxidisers may consume up to 80–90% of the methane produced in certain environments, mitigating its release to the atmosphere (Reeburgh, 2007). The capacity of *Methanoperedens* spp. to accumulate PHA might have implications for the further refinement of these estimates. The inference being that methane would not solely be used for energy production or to increase cell numbers, and that *Methanoperedens* spp. could act as a ‘carbon-capture’ device, especially within carbon-rich anoxic environments whenever they are the main anaerobic methane oxidisers. This would likely be the case for the LFLS test trench, where ammonium, nitrate/nitrite and total dissolved nitrogen are limiting.

## Conclusion

While the *Archaea* inhabiting the LFLS trench water constitute a relatively small portion of its microbial community, they have important roles in biogeochemical cycling, especially with respect to methanogenesis, anaerobic methane oxidation and Fe(III) reduction. The diverse *Methanoperedens* spp. are of special relevance due to their role in methane capture (and subsequent conversion to PHA), Fe cycling, and dissimilatory pathways for nitrate and, possibly, (per)chlorate reduction.

The broad phylogenetic representation of DPANN organisms, which have attracted particular attention in recent years, either for their unusual characteristics or their controversial evolutionary history, is another interesting feature of the LFLS archaeal community. The description and evaluation of the MAGs from lineages LFWA-I to IV constitutes a detailed first examination of several undescribed archaeal lineages.

The proposed LFWA-III lineages, ‘*Ca.* Gugararchaeales’ and ‘*Ca.* Anstonellales’ are amongst the most interesting. Foremost, they are together unusually diverse at LFLS, with 10 out of the 37 MAGs recovered in this study belonging to the two orders, with two families and two genera (and species) newly proposed within each order. Furthermore, they have a cohesive central metabolism that likely spans the entire order with full pathways for the *de novo* biosynthesis of nucleotides, most amino acids, and several vitamins/cofactors. This is duly reflected in the proposed name ‘*Ca.* Gugararchaeum adminiculabundum’, proposed type for LFWA-IIIa, where the specific epithet translates as “self-supporting” relating to the possibility of not needing a symbiotic partner.

## Taxonomic Appendix

Due to several irregularities in the process of naming candidate lineages, the European Nucleotide Archive does not allow certain taxonomic names at taxonomic levels where they should exist. For example, the candidate phylum *Micrarchaeota*, and its defining candidate species ‘*Ca.* Micrarchaeum acidophilum’, have no defined nomenclature at class, order, or family levels. This and other issues related to the nomenclature of uncultivated Bacteria and *Archaea* have been previously discussed in the literature (Whitman, 2016; Konstantinidis et al., 2017, 2019; Oren and Garrity, 2018; Chuvochina et al., 2019; Rosselló-Móra and Whitman, 2019) and we will not provide further discussion on this matter.

Nonetheless, in order to cover these gaps, we feel obliged to suggest several of these intermediate nomenclatural levels, many of which are already covered in the GTDB but not formally proposed in the literature, in addition to the nomenclatural novelties intrinsic to this manuscript.

### Description of ‘*Candidatus* Tiddalikarchaeum’ gen. nov

‘*Candidatus* Tiddalikarchaeum’ (Ti.dda.lik.ar.chae’um. Gunai language, Tiddalik, frog from the Australian Aboriginal mythology; N.L. neut. n. *archaeum*, archaeon, from Gr. adj. *archaios –ê –on*, ancient; N.L. n. neut. *Tiddalikarchaeum*, the archaeon named after the greedy Aboriginal mythological Australian frog that burst with water, referring to the bathtub effect exhibited by the disposal trenches at the Little Forest Legacy Site).

The type species is ‘*Candidatus* Tiddalikarchaeum anstoanum’.

### Description of ‘*Candidatus* Tiddalikarchaeum anstoanum’ sp. nov

‘*Candidatus* Tiddalikarchaeum anstoanum’ (ans.to.a’num. N.L. neut. adj. from ANSTO, Australian Nuclear Science and Technology Organisation, institution managing the Little Forest Legacy Site).

The type material is the metagenome assembled genome (MAG) LFW-252_1 (ERS2655302) recovered from the groundwater of the Little Forest Legacy Site (NSW, Australia). The MAG consists of 1.16 Mbp in 56 contigs with an estimated completeness of 95.7%, redundancy of 1.1%, 16S, 23S, and 5S rRNA gene, and 21 tRNAs. The GC content of this MAG is 36.5%.

### Description of ‘*Candidatus* Tiddalikarchaeaceae’ fam. nov

*Tiddalikarchaeaceae* (Ti.dda.lik.ar.chae.a’ce.ae. N.L. neut. n. *Tiddalikarchaeum* a candidate genus; *-aceae*, ending to denote a family; N.L. fem. pl. n. *Tiddalikarchaeaceae*, the *Tiddalikarchaeum* candidate family).

The family ‘*Candidatus* Tiddalikarchaeaceae’ is circumscribed based on two independent concatenated protein phylogenies of 44 and 91 markers, the latter also supported by the rank normalisation approach as per Parks et al. (2018). The description is the same as that of its sole genus and species. The type genus is ‘*Candidatus* Tiddalikarchaeum’.

### Description of ‘*Candidatus* Tiddalikarchaeales’ ord. nov

‘*Candidatus* Tiddalikarchaeales’ (Ti.dda.lik.ar.chae.a’les. N.L. neut. n. *Tiddalikarchaeum* a candidate genus; *-ales*, ending to denote an order; N.L. fem. pl. n. *Tiddalikarchaeales* the *Tiddalikarchaeum* candidate order).

The order ‘*Candidatus* Tiddalikarchaeales’ is circumscribed based on two independent concatenated protein phylogenies of 44 and 91 markers, the latter also supported by the rank normalisation approach as per Parks et al. (2018). The description is the same as that of its sole genus and species. The type genus is ‘*Candidatus* Tiddalikarchaeum’.

The order is equivalent to CG07-land from Probst, et al. (2018) or o CG07-land in GTDB r89 (Parks et al., 2017).

### Description of “*Nanoarchaeia*” class. nov

“*Nanoarchaeia*” (Na.no.ar.chae’ia. N.L. neut. n. “*Nanoarchaeum*”, a genus; *-ia*, ending to denote a class; N.L. fem. pl. n. “*Nanoarchaeia*”, the “*Nanoarchaeum*” candidate class).

The class “*Nanoarchaeia*” is circumscribed based on two independent concatenated protein phylogenies of 44 and 91 markers, the latter also supported by the rank normalisation approach as per Parks et al. (2018). The description is the same as that of its sole genus and species. The class “*Nanoarchaeia*” is defined as the most inclusive class that includes the genus “*Nanoarchaeum*”. The lineages *Parvarchaeota* (Rinke et al., 2013), *Woesearchaeota* and *Pacearchaeota* (Castelle et al., 2015), and order ‘*Candidatus* Tiddalikarchaeales’ (this work) are contained within “*Nanoarchaeia*”. The type order is “*Nanoarchaeales*” Huber, Auerbach and Podar (2016).

This class is the only class within “*Nanoarchaeota*” Huber & Kreuter (2014), and equivalent to the c Nanoarchaeia in the GTDB r89 (Parks et al., 2017).

### Description of ‘*Candidatus* Gugararchaeum’ gen. nov

‘*Candidatus* Gugararchaeum’ (Gu.ga.rar.chae’um. Dharawal language, *gugara*, kookaburra – bird endemic to Australia, *Dacelo* spp.; N.L. neut. n. *archaeum*, archaeon, from Gr. adj. *archaios –ê –on*, ancient; N.L. neut. n. *Gugararchaeum*, the kookaburra archaeon, honouring the bird, common at the Little Forest Legacy Site, bird emblem of NSW, and typical across Australia).

The type species is ‘*Candidatus* Gugararchaeum adminiculabundum’.

### 3Description of ‘*Candidatus* Gugararchaeum adminiculabundum’ sp. nov

‘*Candidatus* Gugararchaeum adminiculabundum’ (ad.mi.ni.cu.la.bun’dum. L. adj. neut. self-supporting; in reference to the limited external requirements and its suggested independence from a host/symbiote, due to the predicted presence of pathways for the biosynthesis of amino acids, purines, pyrimidines, thiamine, and riboflavin).

The type material is the metagenome assembled genome (MAG) LFW-121_3 (ERS2655302) recovered from the groundwater of the Little Forest Legacy Site (NSW, Australia). The MAG consists of 1.47 Mbp in 76 contigs with an estimated completeness of 96.8%, redundancy of 2.2%, 16S, 23S, and 5S rRNA gene, and 21 tRNAs. The GC content of this MAG is 49.8%.

### Description of ‘*Candidatus* Gugararchaeaceae’ fam. nov

‘*Candidatus* Gugararchaeaceae’ (Gu.ga.rar.chae.a’ce.ae. N.L. neut. n. *Gugararchaeum*, a candidate genus; *-aceae*, ending to denote a family. N.L. fem. pl. n. *Gugararchaeaceae*, the *Gugararchaeum* candidate family).

The family ‘*Candidatus* Gugararchaeaceae’ is circumscribed based on two independent concatenated protein phylogenies of 44 and 91 markers, the latter also supported by the rank normalisation approach as per Parks et al. (2018). The description is the same as that of its sole genus and species. The type genus is ‘*Candidatus* Gugararchaeum’.

### Description of ‘*Candidatus* Burarchaeum’ gen. nov

‘*Candidatus* Burarchaeum’ (Bu.rar.chae’um. Dharawal language, *buru*, kangaroo; N.L. neut. n. *archaeum*, archaeon, from Gr. adj. *archaios –ê –on*, ancient; N. L. neut. n. *Burarchaeum*, an archaeon from the land of the kangaroos, also faunal emblem of Australia).

The type species is ‘*Candidatus* Burarchaeum australiense’.

### Description of ‘*Candidatus* Burarchaeum australiense’ sp. nov

‘*Candidatus* Burarchaeum australiense’ (aus.tra.lien’se. N.L. neut. adj. referring to Australia, country where its first genome was reconstructed).

The type material is the metagenome assembled genome (MAG) LFW-281_7 (ERS2655318) recovered from the groundwater of the Little Forest Legacy Site (NSW, Australia). The MAG consists of 1.20 Mbp in 76 contigs with an estimated completeness of 96.8%, redundancy of 5.4%, 16S and 5S rRNA gene, and 18 tRNAs. The GC content of this MAG is 57.6%.

### Description of ‘*Candidatus* Burarchaeaceae’ fam. nov

‘*Candidatus* Burarchaeaceae’ (Bu.rar.chae.a’ce.ae. N.L. neut. n. *Burarchaeum*, a candidate genus; *-aceae*, ending to denote a family. N.L. fem. pl. n. *Burarchaeaceae*, the *Burarchaeum* candidate family).

The family ‘*Candidatus* Burarchaeaceae’ is circumscribed based on two independent concatenated protein phylogenies of 44 and 91 markers, the latter also supported by the rank normalisation approach as per Parks et al. (2018). The description is the same as that of its sole genus and species. The type genus is ‘*Candidatus* Burarchaeum’.

### Description of ‘*Candidatus* Gugararchaeales’ ord. nov

‘*Candidatus* Gugararchaeales’ (Gu.ga.rar.chae.a’les. N.L. neut. n. *Gugararchaeum*, a candidate genus; *-ales*, ending to denote an order; N.L. fem. pl. n. *Gugararchaeales*, the *Gugararchaeum* candidate order).

The order ‘*Candidatus* Gugararchaeales’ is circumscribed based on two independent concatenated protein phylogenies of 44 and 91 markers, the latter also supported by the rank normalisation approach as per Parks et al. (2018). It constitutes the most inclusive clade that includes the families ‘*Candidatus* Gugararchaeaceae’, and ‘*Candidatus* Burarchaeaceae’. The type genus is ‘*Candidatus* Gugararchaeum’.

The order is equivalent to LFWA-IIIa in this manuscript, and o UBA10214 in the GTDB r89 (Parks et al., 2017).

### Description of ‘*Candidatus* Anstonella’ gen. nov

‘*Candidatus* Anstonella’ (Ans.to.ne’lla. N.L. dim. n. fem. *Anstonella* from ANSTO, Australian Nuclear Science and Technology Organisation, institution managing the Little Forest Legacy Site).

The type species is ‘*Candidatus* Anstonella stagnisolia’

### Description of ‘*Candidatus* Anstonella stagnisolia’ sp. nov

‘*Candidatus* Anstonella stagnisolia’ (s.tag.ni.so’lia. L. v. *stagno* overflow; L. neut. n. *solium - a* tub, bathtub; N.L. adj. fem. *stagnisolia*, overflowing bathtub, in reference to the phenomenon described during heavy rainfalls at the Little Forest Legacy Site trenches).

The type material is the metagenome assembled genome (MAG) LFW-35 (ERS2655287) recovered from the groundwater of the Little Forest Legacy Site (NSW, Australia). The MAG consists of 1.33 Mbp in 68 contigs with an estimated completeness of 97.8%, redundancy of 2.2%, 16S, 23S and 5S rRNA gene, and 21 tRNAs. The GC content of this MAG is 50.3%.

### Description of ‘*Candidatus* Anstonellaceae’ fam. nov

‘*Candidatus* Anstonellaceae’ (Ans.to.nel.la’ce.ae. N.L. neut. n. *Anstonella*, a candidate genus; *-aceae*, ending to denote a family; N.L. fem. pl. n. *Anstonellaceae*, the *Anstonella* candidate family).

The family ‘*Candidatus* Anstonellaceae’ is circumscribed based on two independent concatenated protein phylogenies of 44 and 91 markers, the latter also supported by the rank normalisation approach as per Parks et al. (2018). The description is the same as that of its sole genus and species. The type genus is ‘*Candidatus* Anstonella’.

This family is equivalent to f UBA10161 in the GTDB r89 (Parks et al., 2017).

### Description of ‘*Candidatus* Bilamarchaeum’ gen. nov

‘*Candidatus* Bilamarchaeum’ (Bi.la.mar.chae’um. Dharawal language, *bilama*, freshwater turtle, in reference to their presence still nowadays in the creeks and rivers associated with the Little Forest Legacy Site; N.L. neut. n. *archaeum*, archaeon, from Gr. adj. *archaios –ê –on*, ancient; N.L. neut. n. *Bilamarchaeum*, an archaeon from the turtle lands).

The type species is ‘*Candidatus* Bilamarchaeum dharawalense’.

### Description of ‘*Candidatus* Bilamarchaeum dharawalense’ sp. nov

‘*Candidatus* Bilamarchaeum dharawalense’ (dha.ra.wa.len’se. N.L. neut. adj. pertaining to the Dharawal, traditional owners of the lands where the Little Forest Legacy Site is located).

The type material is the metagenome assembled genome (MAG) LFW-283_2 (ERS2655319) recovered from the groundwater of the Little Forest Legacy Site (NSW, Australia). The MAG consists of 1.27 Mbp in 63 contigs with an estimated completeness of 95.7%, redundancy of 0%, 16S, 23S and 5S rRNA gene, and 20 tRNAs. The GC content of this MAG is 40.0%.

### Description of ‘*Candidatus* Bilamarchaeaceae’ fam. nov

‘*Candidatus* Bilamarchaeaceae’ (Bi.la.mar.chae.a’ce.ae. N.L. neut. n. *Bilamarchaeum*, a candidate genus; *-aceae*, ending to denote a family; N.L. fem. pl. n. *Bilamarchaeaceae*, the *Bilamarchaeum* candidate family).

The family ‘*Candidatus* Bilamarchaeaceae’ is circumscribed based on two independent concatenated protein phylogenies of 44 and 91 markers, the latter also supported by the rank normalisation approach as per Parks et al. (2018). The description is the same as that of its sole genus and species. The type genus is ‘*Candidatus* Bilamarchaeum’.

This family is equivalent to f UBA10214 in the GTDB r89 (Parks et al., 2017).

### Description of ‘*Candidatus* Anstonellales’ ord. nov

‘*Candidatus* Anstonellales’ (Ans.to.nel.la’les. N.L. neut. n. *Anstonella*, a candidate genus; *- ales*, ending to denote an order; N.L. fem. pl. n. *Anstonellales*, the *Anstonella* candidate order).

The order ‘*Candidatus* Anstonellales’ is circumscribed based on two independent concatenated protein phylogenies of 44 and 91 markers, the latter also supported by the rank normalisation approach as per Parks et al. (2018). It constitutes the most inclusive clade that includes the families ‘*Candidatus* Anstonellaceae, and ‘*Candidatus* Bilamarchaeaceae’. The type genus is ‘*Candidatus* Anstonella’.

The order is equivalent to the lineage LFWA-IIIb in this manuscript, and o UBA10214 in the GTDB r89 (Parks et al., 2017).

### Description of ‘*Candidatus* Micrarchaeaceae’ fam. nov

‘*Candidatus* Micrarchaeaceae’ (Mi.crar.chae’a.ce.ae. N.L. neut. n. *Micrarchaeum*, a candidate genus; *-aceae*, ending to denote a family; N.L. fem. pl. n. *Micrarchaeaceae*, the *Micrarchaeum* candidate family).

The family ‘*Candidatus* Micrarchaeaceae’ is circumscribed based on two independent concatenated protein phylogenies of 44 and 91 markers, the latter also supported by the rank normalisation approach as per Parks et al. (2018). It is the most inclusive family that includes the genera ‘*Candidatus* Micrarchaeum’ and ‘*Candidatus* Mancarchaeum’. The type genus is ‘*Candidatus* Micrarchaeum’.

### Description of ‘*Candidatus* Micrarchaeales’ ord. nov

‘*Candidatus* Micrarchaeales’ (Mi.crar.chae.a’les. N.L. neut. n. *Micrarchaeum*, a candidate genus; *-ales*, ending to denote an order; N.L. fem. pl. n. *Micrarchaeales*, the *Micrarchaeum* candidate order).

The order ‘*Candidatus* Micrarchaeales’ is circumscribed based on two independent concatenated protein phylogenies of 44 and 91 markers, the latter also supported by the rank normalisation approach as per Parks et al. (2018). The type genus is ‘*Candidatus* Micrarchaeum’.

The order is equivalent to o Micrarchaeales in the GTDB r89 (Parks et al., 2017).

### Description of ‘*Candidatus* Norongarragalina’ gen. nov

‘*Candidatus* Norongarragalina’ (No.ron.ga.rra.ga.li’na. Dharawal language *Norongarragal*, Dharawal clan group who traditionally occupied the Menai/Lucas Heights area where the present study took place; L. fem. suff., *-ina*, pertaining or belonging to; N.L. dim. n. fem. *Norongarragalina*, the archaeon from the Norongarragal clan).

The type species is ‘*Candidatus* Norongarragalina meridionalis’.

### Description of ‘*Candidatus* Norongarragalina meridionalis’ sp. nov

‘*Candidatus* Norongarragalina meridionalis’ (me.ri.dio.na’lis. L. fem. adj. *meridionalis*, southern; referring to the Southern hemisphere, where its first genome was reconstructed).

The type material is the metagenome assembled genome (MAG) LFW-144_1 (ERS2655293) recovered from the groundwater of the Little Forest Legacy Site (NSW, Australia). The MAG consists of 0.93 Mbp in 49 contigs with an estimated completeness of 93.5%, redundancy of 1.1%, 16S and 5S rRNA gene, and 20 tRNAs. The GC content of this MAG is 57.5%.

### Description of ‘*Candidatus* Norongarragalinaceae’ fam. nov

‘*Candidatus* Norongarragalinaceae’ (No.ron.ga.rra.ga.li.na’ce.ae. N.L. neut. n. *Norongarragalina*, a candidate genus; *-aceae*, ending to denote a family; N.L. fem. pl. n. *Norongarragalinaceae*, the *Norongarragalina* candidate family).

The family ‘*Candidatus* Norongarragalinaceae’ is circumscribed based on two independent concatenated protein phylogenies of 44 and 91 markers, the latter also supported by the rank normalisation approach as per Parks et al. (2018). The description is the same as that of its sole genus and species. The type genus is ‘*Candidatus* Norongarragalina’.

This family is equivalent to f 0-14-0-20-59-11 in the GTDB r89 (Parks et al., 2017).

### Description of ‘*Candidatus* Norongarragalinales’ ord. nov

‘*Candidatus* Norongarragalinales’ (No.ron.ga.rra.ga.li.na’les. N.L. neut. n. *Norongarragalina*, a candidate genus; *-ales*, ending to denote an order, N.L. fem. pl. n. *Norongarragalinales*, the *Norongarragalina* candidate order).

The order ‘*Candidatus* Norongarragalinales’ is circumscribed based on two independent concatenated protein phylogenies of 44 and 91 markers, the latter also supported by the rank normalisation approach as per Parks et al. (2018). The description is the same as that of its sole genus and species. The type genus is ‘*Candidatus* Norongarragalina’.

This order is equivalent to LFWA-II in this manuscript, and o UBA8480 in GTDB r89 (Parks et al., 2017).

### Description of ‘*Candidatus* Micrarchaeia’ class. nov

*Micrarchaeia* (Mi.crar.chae’ia. N.L. neut. n. *Micrarchaeum*, a candidate genus; *-ia*, ending to denote a class; N.L. fem. pl. n. *Micrarchaeia*, the *Micrarchaeum* candidate class.

The class ‘*Candidatus* Micrarchaeia’ is circumscribed based on two independent concatenated protein phylogenies of 44 and 91 markers, the latter also supported by the rank normalisation approach as per Parks et al. (2018). This class is defined as the most inclusive class that includes the orders ‘*Candidatus* Micrarchaeales’, ‘*Candidatus* Gugararchaeales’, ‘*Candidatus* Anstonellales’, and ‘*Candidatus* Norongarragalinales’. The type order is ‘*Candidatus* Micrarchaeales’.

This class constitutes the only class within the phylum ‘*Candidatus* Micrarchaeota’ Baker & Dick (2013). It is equivalent to the c Micrarchaeia in the GTDB r89 (Parks et al., 2017).

### Description of ‘*Candidatus* Methanoperedens vercellense’ sp. nov

‘*Candidatus* Methanoperedens vercellense’ (ver.cel.len’se. L. adj. pertaining or relative to *Vercellae*, Latin name of Vercelli (Italy); in relation to the original MAG code designation, ‘*Ca.* Methanoperedens nitroreducens’ Vercelli, and original location of the sampling from where it was retrieved).

The type material is the metagenome assembled genome (MAG) GCA_900196725 (ERS1800110) recovered from an enrichment culture for anaerobic methanotrophs derived from samples collected from paddy field soil (Vercelli, Italy). The MAG consists of 3.52 Mbp in 250 contigs with an estimated completeness of 96.7%, redundancy of 6.5%, 16S, 23S and 5S rRNA gene, and 21 tRNAs. The GC content of this MAG is 44.1%. Differentiated from other ‘*Ca.* Methanoperedens spp.’ based on ANI and AAI.

### Description of ‘*Candidatus* Methanoperedens batavicum’ sp. nov

‘*Candidatus* Methanoperedens batavicum’ (ba’ta.vi.cum L. adj. *batavicum -us*, relative or pertaining to Batavia (*Ulpia Noviomagus Batavorum*), Latin name of Nijmegen, in reference to the Radboud University Nijmegen, where the initial genome, ‘*Ca.* Methanoperedens nitroreducens’ BLZ1, was studied).

The type material is the metagenome assembled genome (MAG) BLZ1 (GCA_001317315/LKCM00000000.1) recovered from a bioreactor for the enrichment of anaerobic methanotrophs at the Radboud University Nijmegen (Nijmegen, The Netherlands). The MAG consists of 3.74 Mbp in 514 contigs with an estimated completeness of 95.8%, redundancy of 6.6%, 16S, 23S and 5S rRNA gene, and 21 tRNAs. The GC content of this MAG is 40.8%. Differentiated from other ‘*Ca.* Methanoperedens spp.’ based on ANI and AAI.

## Acknowledgements

We acknowledge Shane Ingrey and Raymond Ingrey from the Dharawal Language Program from the Gujaga Fundation for their assistance with the Dharawal language and with their views on aboriginal matters in general. MRW acknowledges the support of the Australian Federal Government NCRIS scheme, via Bioplatforms Australia. MRW and XVC acknowledge support from the New South Wales State Government RAAP scheme and the UNSW RIS scheme. ASK, TEP and TDW acknowledge funding by the Australian Research Council (Discovery Project DP210103727) as well as from the Australian Nuclear Science and Technology Organisation (ANSTO).

## Contribution statement

XVC: Conceptualization, Data curation, Formal analysis, Investigation, Methodology, Project administration, Visualization, Writing – original draft, Writing – review & editing. ASK: Writing – original draft, Writing – review & editing. MWB: Writing – review & editing. MRW: Resources, Supervision, Writing – review & editing. TEP: Resources, Site-specific expertise, Funding acquisition, Project administration. TDW: Resources, Funding acquisition, Project administration, Supervision, Writing – review & editing.

