## Supplementary Information for "Genomic insights into the Archaea inhabiting an Australian radioactive legacy site"

Xabier Vázquez-Campos[1*], Andrew S. Kinsela[2], Mark W. Bligh[2],

Timothy E. Payne[3], Marc R. Wilkins[1] & T. David Waite[2*]

[1] NSW Systems Biology Initiative, School of Biotechnology and Biomolecular Sciences, The University of New South Wales, Sydney, New South Wales 2052, Australia

[2] UNSW Water Research Centre and School of Civil and Environmental Engineering, The University of New South Wales, Sydney, New South Wales 2052, Australia

[3] Environmental Research Theme, Australian Nuclear Science and Technology Organisation, Locked Bag 2001, Kirrawee DC, New South Wales 2232, Australia

* Corresponding authors

### Supplementary Information TOC


Supplementary Information TOC 2

Results and Discussion 3

Carbon metabolism 3

Fixing carbon… or not 4

MAG LFW-68_2 is a versatile protein-degrading *Archaea* 5

LFW-125_1 encodes a respiratory complex I 6

Superoxide reductase in DPANN 7

Filling gaps in the pathways of LFWA-III 7

Biosynthesis of vitamins and cofactors 8

Thiamine biosynthesis 8

Riboflavin biosynthesis 8

Amino acids 9

Lysine and arginine 9

Histidine 10

Cysteine 10

Threonine 10

Serine 11

Aspartate 12

Alanine 12

Tyrosine and phenylalanine 13

Tryptophan 13

Methionine 14

Biosynthesis of nucleosides and nucleotides 15

Glycolysis and gluconeogenesis 17

References 18

### Results and Discussion

#### Carbon metabolism

The central carbon metabolism of the majority of *Archaea* in the LFLS trench waters was either host-dependent glycolytic/fermentative (DPANN) or methane cycling-related (methanogens and AOM). The predominant glycolytic pathway was suggested to be the Embden-Meyerhof-Parnas pathway, based on the lack of 2-keto-3-deoxygluconate kinase (KDG kinase) and 2-keto-3-deoxy-6-phosphogluconate aldolase (KDPG aldolase), along with the absence of key Entner-Doudoroff pathway-enzymes in every MAG, excluding LFW-68_2 (*Thermoplasmata*).

Extracellular CAZymes in the archaeal bins were diverse: a total of 12 different glycoside hydrolase (GH), 9 carbohydrate binding module (CBM), 5 carbohydrate esterase (CE), and 3 polysaccharide lyase (PL) families were detected (Table S13). Proteins containing CBM from the CBM40 and CBM44 families were especially common.

Interestingly, within LFW-68_2, a total of 19 extracellular proteins were predicted to contain a CBM44 domain and 18 of those also contained a polycystic kidney disease (PKD) domain (Table S18). Despite its name, the PKD domains are relatively widespread, mediating protein-protein or protein-carbohydrate interactions in proteins such as collagenases and chitinases, and are also thought to be important components in proteins mediating cell to cell interactions in *Metazoa* (Jing et al., 2002). Only a limited number of CBM44-containing proteins have been well-characterised to date (only 18 sequences, 2 characterised in the CAZy website, 1 August 2019). The CBM44-PKD domain combination has previously been described in the bifunctional cellulase/xyloglucanase of *Clostridium thermocellum* (Najmudin et al., 2006) where it binds cellulose or β1,4-glucans (branched or not, including xyloglucan), and the carbohydrate metabolism protein BT2081 of *Bacteroides thetaiotaomicron* (Yeh et al., 2010). However, it is likely that most of the CBM44-PKD containing proteins (12 out of the 18) are acting as proteases rather than on carbohydrates, given their similarity with proteases in the MEROPS database, mainly M09B (Table S18). Further details into the detritivorous/proteolytic lifestyle of LFW-68_2 are explored in its own section.

#### Fixing carbon… or not

To date, there are six known pathways for the autotrophic fixation of $CO_2$, yet the Calvin-Benson-Bassham (CBB) cycle, with RuBisCO as key enzyme, remains the most widespread pathway (Lannes et al., 2019). RuBisCO and RuBisCO-like proteins are currently classified based on their phylogeny into five clades named I to IV, in addition to the II/III mixed type (Wrighton et al., 2016).

Either RuBisCO form II/III, III or IV (RuBisCO-like proteins) were detected in 22 of the 37 MAGs (Figure 2, Table S10). Despite their names, these types of RuBisCO are not necessarily linked with self-sufficient carbon fixation through the CBB cycle.

Wrighton et al. (2016) demonstrated that the RuBisCO type II/III from *Peregrinibacteria* (PER) (and potentially others) can support autotrophic growth on its own, although it is not clear that it would function as in the original organism, nor that it can be generalised to every RuBisCO form II/III. The *Pacearchaeota* LFW-170_1 and LFW-170_3, as well as the LFW-283_2 MAG from LFWA-III are the only *Archaea* harbouring this form of RuBisCO.

On the other hand, the RuBisCO form III, was present in 19 MAGs, and has been shown to be involved in AMP metabolism and not related to the CBB cycle (Sato et al., 2007). Although it was demonstrated that form III does fix carbon and could, to some extent, support growth (Sato et al., 2007), it has been postulated that it would be insufficient to sustain all C-requirements of the cell (Sato et al., 2007). The *Archaea* containing RuBisCO form III, often rely on the fixation of carbon through alternate pathways, e.g. reductive acetyl-CoA pathway (also known as Wood Ljungdahl pathway). Indeed, the acetyl-CoA decarbonylase/synthase, a key protein complex of the reductive acetyl-CoA pathway, was detected in all methanogen and methane oxidiser MAGs, at least partially. Thus, the reductive acetyl-CoA pathway would appear to be the main C fixation mechanism in the LFLS *Archaea*.

#### MAG LFW-68_2 is a versatile protein-degrading *Archaea*

In settings where assimilable nitrogen can be scarce, as is the case at LFLS, being able to fix $N_2$ can be advantageous. Dinitrogenase components I and II were only detected in the *Methanoperedens* spp. MAGs with ≥90%C and within a single methanogen (LFW-151_2; Figure 2 and Table S10). While the dinitrogenase in LFW-151_2 can be easily defined as [MoFe]-nitrogenase, the nitrogenases of the *Methanoperedens* spp. MAGs do not match any specific type based on the metal centre. This is concordant with previous work on the metal centre of nitrogenases where the anaerobic methane oxidising *Archaea* were classified as "undefined" type nitrogenases (Lloyd et al., 2013) and likely belonging to the Nif-D lineage (Laskar et al., 2011).

However, nitrogen fixation is an energetically costly process. Sediments accumulate detrital organic matter that can provide a source of carbon and nitrogen. Some archaeal lineages such as *Thorarchaeota* (Seitz et al., 2016) and MBG-D (Marine Benthic Group D) (Lloyd et al., 2013) are well equipped to exploit detrital material. Similar to the related MBG-D archaea SCGC AB-539-N05 (ALXL00000000.1) and SCGC AB-539-C06 (AOSH00000000.1) (Lloyd et al., 2013), LFW-68_2 (*Thermoplasmata*) possess an expanded repertoire of peptidases. With 59.84 MEROPS matches per Mbp (compared to ~50/Mbp for the aforementioned MBG-D and 25-35/Mbp for most other archaea), LFW-68_2 is the archaeal genome with the highest density of peptidase-coding genes (Figure S1). The collection of peptidases in the genome of LFW-68_2 is especially expanded in C25.001 cysteine peptidases (15 copies), M09.002 metallopeptidases (17 copies), and M08.020 metallopeptidases (8 copies). A total of 26 extracellular peptidases provides LFW-68_2 with an exhaustive machinery for the degradation of detrital proteins.

The proteolytic capabilities of LFW-68_2 are even more exceptional than just a broad repertoire of proteases. We predicted an extracellular protease from the S12 family (PID 281199). The family S12, along with S11 and S13, constitute the SE clan which includes peptidases with specialised roles in the bacterial cell-wall metabolism; i.e. they act on D-amino acids (Laskar et al., 2011). A cytoplasmic M19 dipeptidase (PID 342444) was also predicted and is one of the few dipeptidases able to cleave dipeptides regardless of whether the C-terminal residue is a D- or L- isomer (Hooper, 2013). Lastly, we predicted a putative operon (PID 515476-85, Figure S8B) containing homologues to the peptide-binding substrate binding protein-dependent ABC transport system (Dpp/Ddp/Opp/App/Gsi), used for the import of D,D-amino acids (Maqbool et al., 2011). In contrast to other well-known examples where the operons are limited to a single gene per protein in the transport system, the organisation of the peptide-binding transport operon in LFW-68_2, i.e. DdpFDCBAFFCBA (Figure S8B), suggests that a duplication/rearrangement/recombination event occurred somewhere during its evolutionary history. The above three points suggest that LFW-68_2 might not have a role in the degradation of proteins but in the degradation of D-amino acid-containing organic matter.

The D-amino acid containing necromass is one of the most recalcitrant components, often found in bacterial cell walls, as well as enriched in aged sediments, as a product of amino acid racemisation (Lomstein et al., 2012; Lloyd et al., 2013). The presence of the three aforementioned components (i.e. extracellular protease S12, M19 dipeptidase and oligo-/di-peptide transport system) support the hypothesis that LFW-68_2 is a detritivorous *Archaea* able to feed on bacterial cell walls and recalcitrant biomass.

While all other MAGs have some kind of amino acid transporters (MFS and/or ABC), the transporter systems for oligopeptides are mainly limited to App (min 4 aa chains) in *Methanomicrobia*.

While the *Nitrosotalea*-related MAG LFW-283_4_5 would be expected to contribute to the nitrogen cycle as ammonia oxidising archaea, the recovered bin is partial, so we were not able to predict any proteins involved in this process (Table S10). No other MAG showed any indication of dissimilatory pathways for inorganic nitrogen compounds.

No evidence of sulfur dissimilatory pathways was found in any of the recovered genomes.

### LFW-125_1 encodes a respiratory complex I

Like most DPANN, the DPANN MAGs at LFLS lack traces of any typical electron transfer chain aside from ATPase. The only exception is LFW-125_1 (*Micrarchaeales*), more related to ARMAN-2 than to other DPANN in the trenches. This archaeon encodes a respiratory complex I, mostly in a single operon (NuoABCDHIJKLN) with NuoM encoded in a different contig. While most of the components can be identified based on arCOGs, NuoA, NuoJ and NuoK fall below thresholds for both arCOGs, Kofam and lack any specific feature identifiable through InterproScan (Table S19). In addition, only NuoB and NuoN are properly identified by KofamScan.

Another interesting point is that this respiratory complex I lacks a G subunit typical of most "modern" complex I, and, with only 11 components, it resembles the proposed last common ancestor of the complex I proposed by Moparthi and Hägerhäll (2011).

#### Superoxide reductase in DPANN

The presence of superoxide reductases in LFW-252_1 and other LFLS DPANN genomes, 6 out of the 25 DPANN (a few *Pacearchaeota* and LFW-252_1), is quite unusual, especially when compared with extant annotations, e.g. AnnoTree (r89), where only 3/151 DPANN genomes contain SOR. While SOR-containing genomes have been typically described as strict anaerobes, there are multiple recent examples that contradict this behaviour. Bacteria such as *Sulfurimonas autotrophica* (Inagaki et al., 2003), *Halarcobacter bivalviorum* (Levican et al., 2012), *Escherichia albertii* (Huys et al., 2003), and *Sulfurovum lithotrophicum* (Inagaki et al., 2004), have all been shown to use SOR for the detoxification of superoxide, despite none of them being classified as strict anaerobes. However, it is also true that these organisms utilising SOR prefer to grow under aerobic conditions with oxygen concentrations below atmospheric partial pressures.

#### Filling gaps in the pathways of LFWA-III

During the examination of the metabolic reconstruction of the individual archaeal genomes, it was observed that some DPANN from the LFWA-III lineage, exemplified by LFW-121_3, possess nearly complete pathways for the synthesis of amino acids, purine and pyrimidine nucleotides, riboflavin, and thiamine (Figure 4). However, several of the LFW-121_3 pathways, and LFWA-III in general, had a limited number of gaps. Manual curation with predicted functions from other databases combined with literature searches was able to fill some of those gaps or, at least, provide reasonable candidate proteins that may carry on those functions. These are discussed in the sections below.

#### Biosynthesis of vitamins and cofactors

##### Thiamine biosynthesis

The biosynthesis of thiamine arises from the condensation of a compound with a thiazole ring and an aminopyrimidine derivative. In the extant model in *Archaea*, based on *Euryarchaeota* and *Crenarchaeota*, the final aminopyrimidine derivative, HMP-PP (4-amino-5-hydroxymethyl-2-methylpyrimidine diphosphate) derives from a metabolite from the purine/histidine pathways, AIR (5-amino-1-(5-phospho-β-D-ribosyl)imidazole), while the thiazole ring is contained in THZ-P (4-methyl-5-(β-hydroxyethyl)thiazolium phosphate) derived from glycine and NAD (Hwang et al., 2017; Maupin-Furlow, 2018).

According to the KEGG modules, LFW-121_3 possesses all the enzymes required for the biosynthesis of thiamine. However, careful examination shows the absence of the enzymes in the thiazole branch leading to the *de novo* formation of a thiazole ring. While the enzyme responsible for the second step in *Archaea* is still a mystery, the first one is present in multiple *Archaea* (Maupin-Furlow, 2018). LFW-121_3 can still synthesise thiamine by salvaging THZ, which is phosphorylated to THZ-P by ThiM.

##### Riboflavin biosynthesis

The pathway for the biosynthesis of riboflavin is nearly complete in LFW-121_3. The phosphatase required for the cleavage of the phosphate group from 5-Amino-6-(5'-phospho-D-ribitylamino)uracil, EC3.1.3.104, is absent. However, no candidate enzyme has been proposed that could fill this gap in *Archaea* (Rodionova et al., 2017).

In Bacteria, this step is often carried out by orthologues of YcsE (*Bacillus subtilis*, COG0561) or YigB/YbjI (*Escherichia coli*, COG1011). Despite belonging to different orthologous groups, they are both members of the haloacid dehalogenase enzyme superfamily (HAD). Like most HAD enzymes, HAD phosphatases are characterised by their considerable substrate promiscuity (Haase et al., 2013; Huang et al., 2015). Indeed, Cof-like phosphatases, a common family of HAD phosphatases, have, for example, 7 paralogues with overlapping substrate specificity in *E. coli* (Mendonça et al., 2011). In contrast, the endosymbiotic bacterium '*Ca.* Blochmannia pennsylvanicus' is capable of utilising 4 of the same 5 substrates with just a single Cof-like phosphatase (Mendonça et al., 2011).

The ubiquity of biochemical reactions driven by phosphatases and the existence of a linear correlation between genome size and the number of phosphatase annotations (Figure S9) in addition to their rather redundant functionality, suggests that in case of a genome reduction, phosphatase-encoding genes could be lost without a reduction of function as far as they can be supplemented by a related promiscuous phosphatase, as in the case of '*Ca.* Blochmannia pennsylvanicus'.

#### Amino acids

L-leucine, L-valine biosynthesis pathways in LFW-121_3 are predicted by both Pathway Tools and KEGG. However, many other amino acid biosynthesis pathways show gaps with either one or both methods, conflicting annotations, etc. These pathways are discussed further in this section.

##### Lysine and arginine

It has been proposed that leucine, lysine and arginine pathways share a common ancestor given the numerous enzymes from the different paths that belong to the same orthologous groups (Fondi et al., 2007).

Lysine is produced via L-2-aminoadipate in LFW-121_3. The genes in the biosynthesis of lysine are encoded in two separate gene clusters: *lysYZJK-argCB* (PID 405578-83) and *lysWX* (PID 589123-4). Based on phylogenetic analysis (Figure S10), the predicted LysJ ([LysW]-aminoadipate semialdehyde transaminase) is closely related to the ArgD (acetylornithine aminotransferase) in *Thermus thermophilus* (Horie et al., 2009) and *Leptospira interrogans*, known to also function as LysJ in the lysine biosynthesis pathway via L-2-aminoadipate. Bifunctional LysJ/ArgD homologues are also known in other *Archaea* such as *Thermococcus kodakarensis* (Yoshida et al., 2016) and *Sulfolobus acidocaldarius* (Ouchi et al., 2013) where most proteins involved in the biosynthesis of Lys and Arg are shared between both pathways. A screening on the complementarity of the diverse aminotransferases from the Arg and Lys biosynthetic pathways in *Escherichia coli* showed that all but three enzymes were intrinsically promiscuous (Lal et al., 2014). All this suggests that the enzyme in LFW-121_3 may have the same bifunctionality.

The presence of ArgJ (PID 148337) indicates that the biosynthesis of arginine follows the L-arginine biosynthesis II (acetyl cycle) pathway (MetaCyc nomenclature). While the canonical ArgA is missing, ArgJ is known to be bifunctional, functioning also as ArgA in many organisms (Marc et al., 2000; Xu et al., 2007), allowing closure of another pathway gap.

##### Histidine

According to Pathway Tools, the L-histidine biosynthetic pathway is complete but KEGG fails to identify HisB (histidinol phosphatase, EC3.1.3.15). The HisB identified by Pathway Tools (PID 474067) cannot be recognised as such by HAMAP (MF_00076) although it still belongs to the same protein family of the canonical HisB based on CDD (cd07432, PHP_HisPPase). A different HisB, annotated by EggNOG (PID 474067), matches the required HAMAP profile with greater ease, providing a better candidate protein.

##### Cysteine

Cysteine is produced in LFW-121_3 from Ser with O-acetyl-L-serine as an intermediate (Makino et al., 2016), and driven by CysE (TIGR01172, PID 405660) and CysK (TIGR01139, PID 405659), detected by TIGRFAM and KEGG, but overlooked by Pathway Tools.

##### Threonine

The biosynthesis of L-threonine requires homoserine kinase (HSK, ThrB) for its synthesis from aspartate semialdehyde. However, no canonical HSK was found in the genome of LFW-121_3. Instead, there is an alternate ApgM-like (2,3-bisphosphoglycerate-independent phosphoglycerate mutase) homoserine kinase (TIGR02535, PID 405464). This alternate HSK seems to be common in *Archaea*, being present in 48 of the 279 genomes screened compared to 88 containing the canonical forms of HSK (TIGR00191 or TIGR00938). The alternate HSK has also been proposed to be involved in the biosynthesis of Thr in other non-model organisms such as the bacterium *Dehalobacter restrictus* (Rupakula et al., 2015).

#### Serine

The three main steps characteristic for the biosynthesis of Ser are catalysed, in order, by SerA, SerC and SerB. While a canonical SerA (D-3-phosphoglycerate dehydrogenase / 2-oxoglutarate reductase) is available, the specific enzymes for the next steps are missing in the genome of LFW-121_3. SerA catalyses the formation of 3P-hydroxypyruvate (3PHP) from 3P-D-glycerate. The next reaction is the transamination of 3PHP to O-phosphoserine by SerC (phosphoserine aminotransferase). However, its absence in a serine-synthesising *Archaea* is not particularly novel. In the genome of LFW-121_3 we find a protein candidate (PID 1010399), with some ambiguous annotations that might suggest its role as SerC, e.g. PIRSF000524/IPR024169 (serine-pyruvate aminotransferase/2-aminoethylphosphonate-pyruvate transaminase). The closest matches at UniProtKB/Swiss-Prot to PID 1010399 (36.2% identity), is MJ0959 from *Methanocaldococcus jannaschii*. Both MJ0959 from *M. jannaschii*, and the homologous MMP0391 from *Methanococcus maripaludis*, exhibit bifunctionality as both aspartate aminotransferase (AspC) and SerC (Helgadóttir et al., 2007). Protein 1010399 has a similar size (355 aa vs 385 aa in MJ0959 and 382 aa in MMP0391), same orthology (arCOG00082) and the same domain annotations (i.e. Pfam, PIRSF, Gene3D and SUPFAM). Hitherto, we suggest that PID 1010399 may function in a similar way, despite the rather low sequence identity.

The last step of the Ser biosynthesis pathway constitutes the hydrolysis of the phosphate group in phosphoserine. This process, carried out by SerB, is probably the most understudied in the biosynthesis of Ser. The canonical phosphoserine phosphatase (PSP) belongs to the HAD hydrolase superfamily and the broad orthologous group COG0560. However, several alternate PSP have been discovered in recent years. In *Hydrogenobacter thermophilus*, the PSP activity is catalysed by a cofactor-dependent phosphoglycerate mutase (dPGM)-like protein from the histidine phosphatase family (COG0406, PF00300) likely to be broadly distributed amongst *Firmicutes* and *Cyanobacteria* (Chiba et al., 2012). In *Thermus thermophilus*, the PSP activity is also performed by a member of the HAD hydrolase superfamily (TtPSP, COG1011, TIGR01549+TIGR01509), but more closely relate to the 3-phosphoglycerate dehydrogenase (PGDH) than to the canonical PSP (Chiba et al., 2019). The predicted proteome of LFW-121_3 contains at least two instances of TtPSP-like proteins matching the orthology and the dual TIGRFAM domain annotation, PID 589042 and PID 296056.

##### Aspartate

L-aspartate is synthesised through the oxaloacetate produced by action of the unusual phosphoenolpyruvate carboxykinase (PID 405563, PF01293, PEPCK, below threshold as arCOG06073) that contains a STAS regulatory domain (PS50801) towards the C-terminus. The STAS domains have been often associated with their role as anti-anti-sigma factors (Sharma et al., 2011). Although more recent studies challenge the idea of STAS domains as anti-anti-sigma factors as their only possible function (Thompson and Visick, 2015), it seems that their involvement in regulatory processes is still a common denominator.

##### Alanine

Alanine can be produced in two ways, either by the transamination of pyruvate using Glu as amino donor or as a byproduct of the desulfurisation of Cys to provide sulfur for the formation of [Fe-S] clusters and other S-containing molecules (Frazzon and Dean, 2003). In either case, KEGG predictions failed to detect the required enzymes, and Pathway Tools only provides a suggestion for the second path. A close look into the EggNOG and InterproScan results indicate that LFW-121_3 has, indeed, the enzymes needed for the biosynthesis of Ala. For the transamination reaction there is one alanine aminotransferase (arCOG01131, PID 1198997). For the desulfurisation path, there are two candidate cysteine desulfurase (arCOG00065; PID 1267536 and PID 1267551).

#### Tyrosine and phenylalanine

Both tyrosine and phenylalanine are synthesised via a prephenate intermediate. In the case of Tyr, prephenate is first transformed into 4-hydroxyphenylpyruvate by prephenate dehydrogenase (arCOG00245, PF02153; PID 405403). In the biosynthesis of Phe, prephenate is instead converted to phenylpyruvate by prephenate dehydratase (arCOG00255, PF00800, PS51171; PID 1104438). In both cases, the final reaction is a transamination which, depending on the organism, can come in many shapes. In our case, HisC would be the responsible enzyme for the transference of the amine group from Glu to phenylpyruvate and 4-hydroxyphenylpyruvate. Mutants of *Bacillus subtilis* lacking either of the terminal aminotransferases from the Phe and Tyr retain prototrophy for both amino acids as a result of HisC activity (Nester and Montoya, 1976; Weigent and Nester, 1976). Similar enzyme promiscuity has been described in *Thermotoga maritima* (Fernandez et al., 2004) and *Corynebacterium glutamicum* (Marienhagen et al., 2008). In addition, HisC is one of the only two proteins from the histidine biosynthetic pathway predicted to be absent from LUCA and suggested to evolve from ‘generic’ aminotransferases in which LUCA would have relied upon (Fondi et al., 2009).

#### Tryptophan

Both KEGG and Pathway Tools fail to provide a complete pathway for the biosynthesis of Trp. In the case of KEGG, the examination of the pathway modules suggests two missing key proteins: TrpG and TrpF.

The TrpG (glutamine amidotransferase subunit of anthranilate synthase) is apparently also missing from the EggNOG annotation. On closer examination, and combining the annotations from InterProScan, TrpG is clearly represented by PID 148344 (arCOG00086, TIGR00566, PR00097, cd01743) but erroneously labelled as TrpD by EggNOG. The actual TrpD is PID 1104434 (arCOG02012, TIGR01245, MF_00211).

Despite possessing all the other enzymes for the biosynthesis of Trp, the canonical phosphoribosyl anthranilate (PRA) isomerase (TrpF) is not present amongst the predicted proteins of LFW-121_3. All domain database annotations fail to recognise any typical signature for PRA isomerase domains, e.g.

PF00697 (PRAI domain), cd00405, MF_00135, or TIGR00566, or even the alternate forms such as those defined by TIGR01919 (bifunctional *hisA*-*trpF*). However, the PRAI activity is not necessarily exclusive of a PRAI domain, and, in general, it seems that enzymes acting on phosphoribosylated substrates often have overlapping substrate activities (Patrick and Matsumura, 2008). For example, the PurF (glutamine phosphoribosylpyrophosphate amidotransferase) from *E. coli* is known to be able to cover the nutritional requirements of the bacterium in the absence of TrpF (Patrick and Matsumura, 2008). Another isomerase involved in the biosynthesis of amino acids, HisA, is a known homologue of TrpF (Due et al., 2011; Plach et al., 2016; Newton et al., 2017). Organisms that have lost their TrpF may carry a bifunctional homologue of HisA, known as PriA, that covers the activities of both HisA and TrpF. Even if PriA is not present, several HisA are known to be able to process TrpF substrate with enough affinity to allow growth (Plach et al., 2016). Artificial evolution experiments have demonstrated that the *N*′-[(5′-phosphoribosyl)formimino]-5-aminoimidazole-4-carboxamide ribonucleotide (ProFAR) isomerase (HisA) in *Salmonella enterica* can retain ProFAR isomerase activity and gain low level PRA isomerase activity enough to support growth with only two mutations (Näsvall et al., 2012), yet not change its domain profile making it virtually impossible to detect such changes based on sequence alone (Table S20).

#### Methionine

To date, a minimum of 18 different methionine biosynthesis pathways have been proposed, all originating from homoserine as precursor (Gophna et al., 2005). The biosynthesis of methionine is usually defined in three phases: i) homoserine (HS) activation by acylation, forming *O*-acetyl- or *O*-succinyl-homoserine (OAHS or OSHS); ii) sulfur incorporation to produce homocysteine (Hcy) by MetY or MetB; and, iii) methylation of homocysteine to Met. Like most organisms, we can easily identify the proteins responsible for steps i) (homoserine *O*-acetyltransferase, MetX; PID 148362) and iii) (cobalamin-independent methionine synthetase, MetE; PID 1101347-8) in LFWA-121_3. However, the incorporation of sulfur appears to be a cryptic step in many *Archaea* and *Bacteria*. For example, *Archaeoglobus fulgidus*, '*Ca.* Caldiarchaeum subterraneum', *Lokiarchaeum* sp. GC14_75 and *Korarchaeum cryptofilum*, all contain the required enzymes to produce OAHS (or at least HS), as well as methionine from Hcy but they lack of any known candidate to perform the sulfur incorporation (KEGG, Jan 2019). Nonetheless, two of the other highly complete LFWA-III genomes do contain genes encoding for enzymes able to catalyse the sulfur incorporation onto the *O*-acylated homoserine: LFW-281_6_1 has a MetB (PID 208256, arCOG00060), while LFW-281_7 has a MetB (PID 1221042) and a MetY (arCOG00061, PID 666916). In any case, the lack of any kind of cystathionine β-synthase in LFW-121_3 or in any of the LFWA-III members, indicates that Met biosynthesis would be through the direct incorporation of sulfur into *O*-acetylhomoserine and not through a cystathionine intermediate.

#### Biosynthesis of nucleosides and nucleotides

The pathways for the *de novo* biosynthesis of purine and pyrimidine nucleosides and nucleotides are almost perfectly reconstructed in LFW-121_3. The apparent gaps in the metabolic models align in most cases to known knowledge gaps regarding the metabolism of *Archaea*.

In the case of the pyrimidine nucleotide biosynthesis pathway, both Pathway Tools and KEGG fail to predict one step on each. However, missing predictions are not the same for both tools, generating a complete pathway when combined.

The biosynthesis of purine nucleotides seems to have three missing enzymes: the 5-(carboxyamino)imidazole ribonucleotide synthase (PurK, $N^5$-CAIR synthase, EC6.3.4.18), the PurH/PurO, and the guanylate kinase (GK).

The PurK protein is involved in an intermediate step for the biosynthesis of inosine 5'-monophosphate (IMP), a required precursor for the formation of purine nucleotides. The said step is the production of CAIR (5-amino-1-(5-phospho-D-ribosyl)imidazole-4-carboxylate) from AIR. In most Eukarya, this is often performed in a single reaction catalysed by a class II PurE (PurE-II, AIR carboxylase, EC4.1.1.21) using $CO_2$ as substrate (Armenta-Medina et al., 2014). An alternate process, present in most *Bacteria*, *Archaea* and some *Eukarya* lineages, requires two steps: first, PurK catalyses the formation of the $N^5$-CAIR intermediate using $HCO_3^-$ instead of $CO_2$ as substrate and, second, a $N^5$-CAIR mutase (class I PurE, PurE-I, EC5.4.99.18), evolutionarily related to the AIR carboxylase, converts $N^5$-CAIR in CAIR

(Armenta-Medina et al., 2014). However, the PurE-I containing organisms do not necessarily need a copy of PurK. The carboxylation of AIR to $N^5$-CAIR can actually occur spontaneously under environmental conditions with high concentrations of $HCO_3^-$ (Mathews et al., 1999; Meyer et al., 1999; Armenta-Medina et al., 2014). LFW-121_3's genome, in a similar manner to *Archaeoglobus fulgidus*, encodes a PurE-I (PID 58321) but not a PurK.

The last step in the *de novo* biosynthesis of purine nucleotides is the cyclation of the FAICAR (5-formamido-1-(5-phospho-D-ribosyl)-imidazole-4-carboxamide) moiety that will result in the formation of the hypoxanthine in IMP. This process is performed by enzymes with the IMP cyclohydrolase (IMPCH) activity. In most Bacteria and Eukarya, the protein PurH carries the IMPCH domain as well as the 5-amino-1-(5-phospho-D-ribosyl)imidazole-4-carboxamide (AICAR) formyl transferase (AICARFT) (Costa Brandão Cruz et al., 2019). In *Archaea*, however, IMPCH and AICARFT activities are often separated. The IMPCH activity is carried out by PurJ or PurO, and AICARFT by PurV or PurP. While no protein with IMPCH activity was predicted in the genome of LFW-121_3, several other LFA-III *Archaea* present a typical PurH in conjunction with two different PurP proteins (LFW-29, LFW-35, LFW-281_3_2, LFW-281_5_4, LFW-281_7 and LFW-283_2). Given that the PurH is present in all other LFA-III genomes with high completeness, we suggest that its absence in LFW-121_3 is likely due to be missing in the assembly rather than actually lacking the *purH* gene. The presence of PurH along with two different PurP, is akin to the reported for *Methanosarcinaceae* and *Thermoplasmatales* archaeon BRNA1 by Costa Brandão Cruz *et al.* (2019).

Guanylate kinase (GK) activity (conversion of GMP to GDP) has been detected in *Archaea*. However, the responsible enzyme remains elusive (Armenta-Medina et al., 2014). It has been suggested that given the high similarities with other nucleoside monophosphate (NMP) kinases, especially adenylate kinase (AK), and the minimal changes required to shift the specificity of a guanylate kinase enzyme, it is possible that the actual GK in *Archaea* might be being labelled as AK instead (Stolworthy and Black, 2001; Armenta-Medina et al., 2014). The genome of LFW-121_3, as in many other *Archaea*, contains two possible AK: PID 1267582 and PID 474048. While the first is a canonical archaeal AK (MF_00234, arCOG01039), the second is related to the AK isozyme 6 of eukaryotic AK, and defined as putative adenylate kinase by HAMAP given the lack of experimental evidence (MF_00039, arCOG01038). The EggNOG database attributes a rather different annotation based on the broad specificity of these enzymes, i.e. NMP kinases (EC2.7.4.4). Based on this, PID 474048 could not only perform as an as GK but as reversible IMP kinase too, filling an additional gap needed for the salvage of ITP.

#### Glycolysis and gluconeogenesis

Unlike the neighbouring *Micrarchaeota*, LFWA-III utilises the Embden–Meyerhof–Parnas (EMP) pathway instead of the Entner–Doudoroff pathway, based on the absence of KDG kinase and KDPG aldolase. Based on both KEGG and Pathway Tools, the EMP pathway would be complete but missing one key enzyme commonly absent in other *Archaea*, the phosphofructokinase (PFK).

No canonical PFK, or any known variations (i.e. alternate phosphate donors other than ATP), was found in any of the DPANN genomes analysed, at least not when using type-specific profiles (TIGR03828, TIGR02477, TIGR02045, TIGR02482, TIGR02483, MF_01979, PIRSF036482). A search for other, ambiguous, annotations revealed the presence of ADP-specific phosphofructokinase/glucokinase (ADP-PFK/GK) domains (PS51255, PF04587, IPR007666) in proteins of a number of LFWA-III genomes and other *Archaea*. In every case, the proteins belong to arCOG03371 or, at least, would have the best orthologous match in this group. The ADP-PFK/GK proteins matching arCOG03371 had the best match on the bifunctional ADP-PFK/GK of *Methanothrix sohngenii* (F4BV59, syn. *Methanosaeta concilii*) or *Methanocaldococcus jannaschii* (Q58999) (Sakuraba et al., 2002). The lack of genes coding for enzymes with specific PFK activity is not unusual in *Archaea* even when there is a complete gluconeogenesis pathway and/or total absence of the ED pathway. The fact that two of the few model *Archaea* fit the situation described here-in suggests that the bifunctional PFK/GK enzymes are rather common among *Archaea* with the EMP pathway.
