## Supplementary figures and images for "Genomic insights into the Archaea inhabiting an Australian radioactive legacy site"

### Figure S1

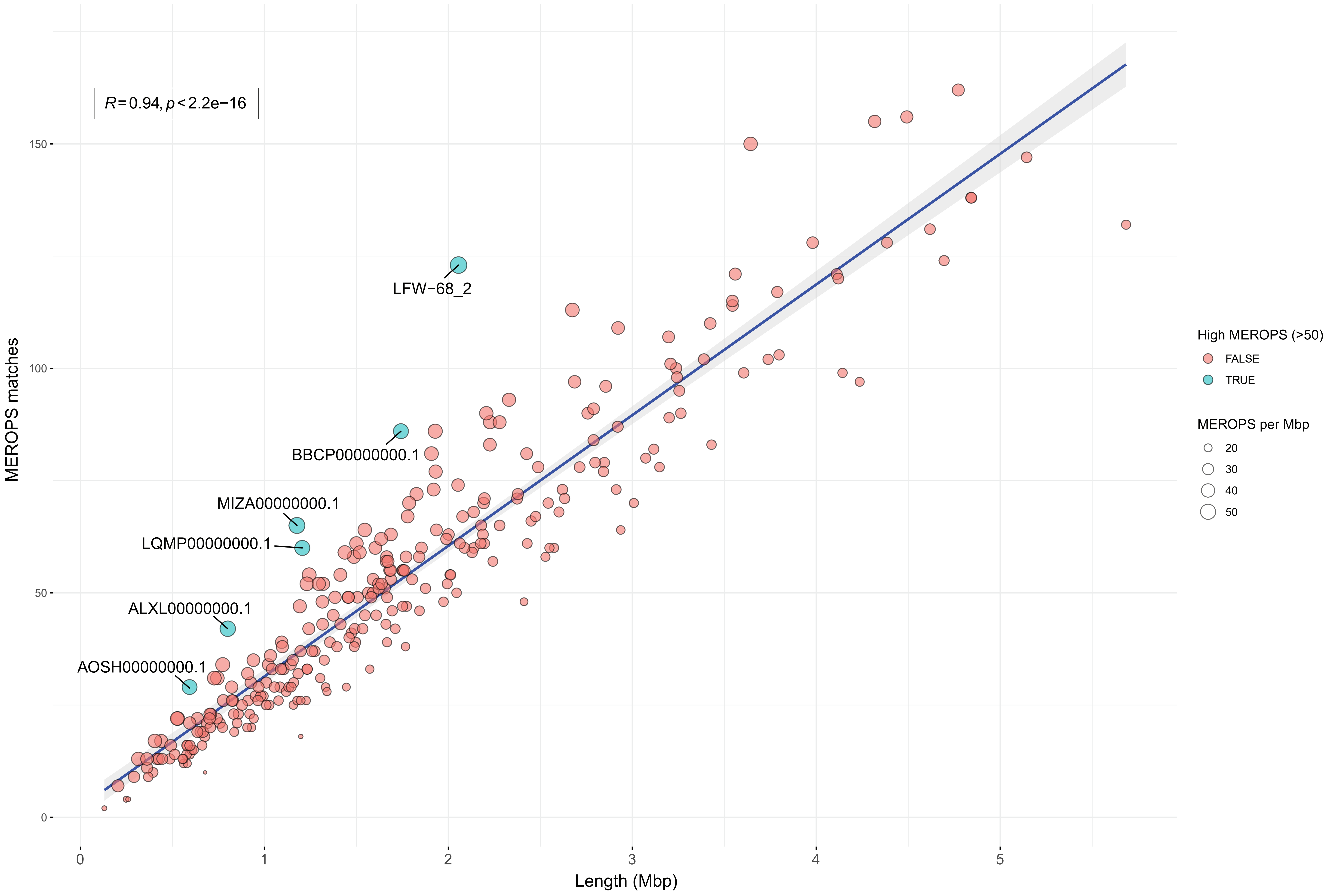

### Figure S2

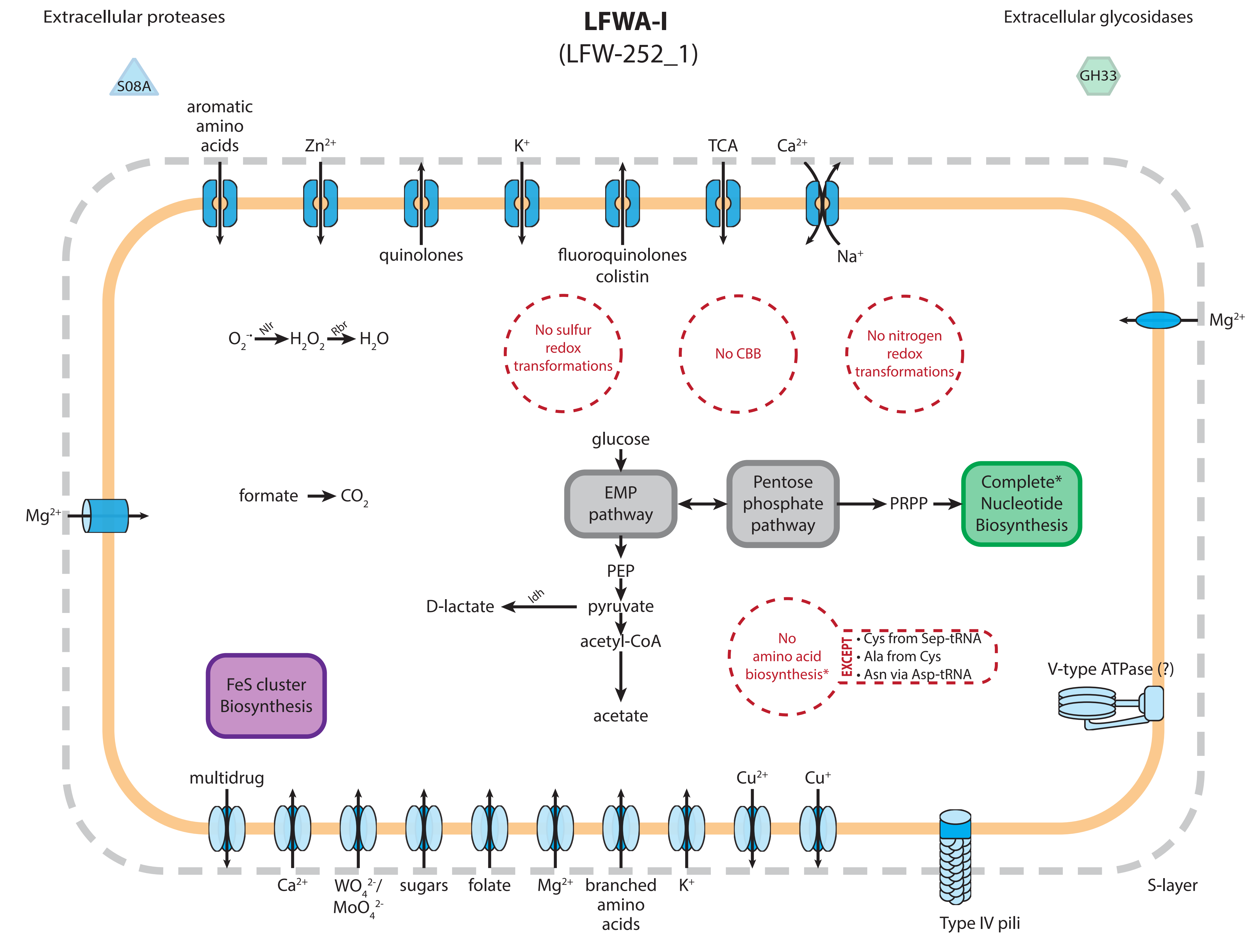

### Figure S3

#### LFWA-II
(LFW-144_1)

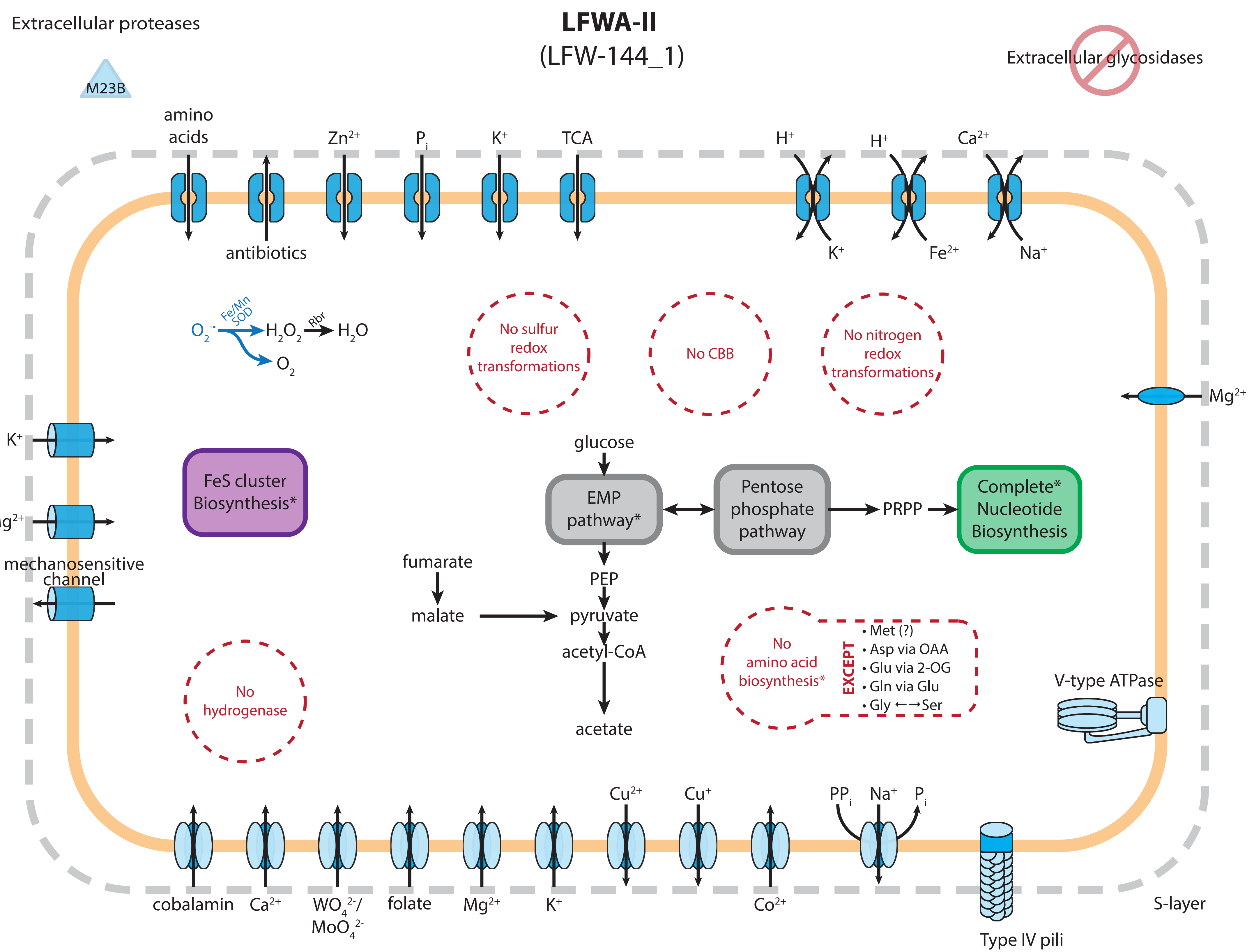

### Figure S4

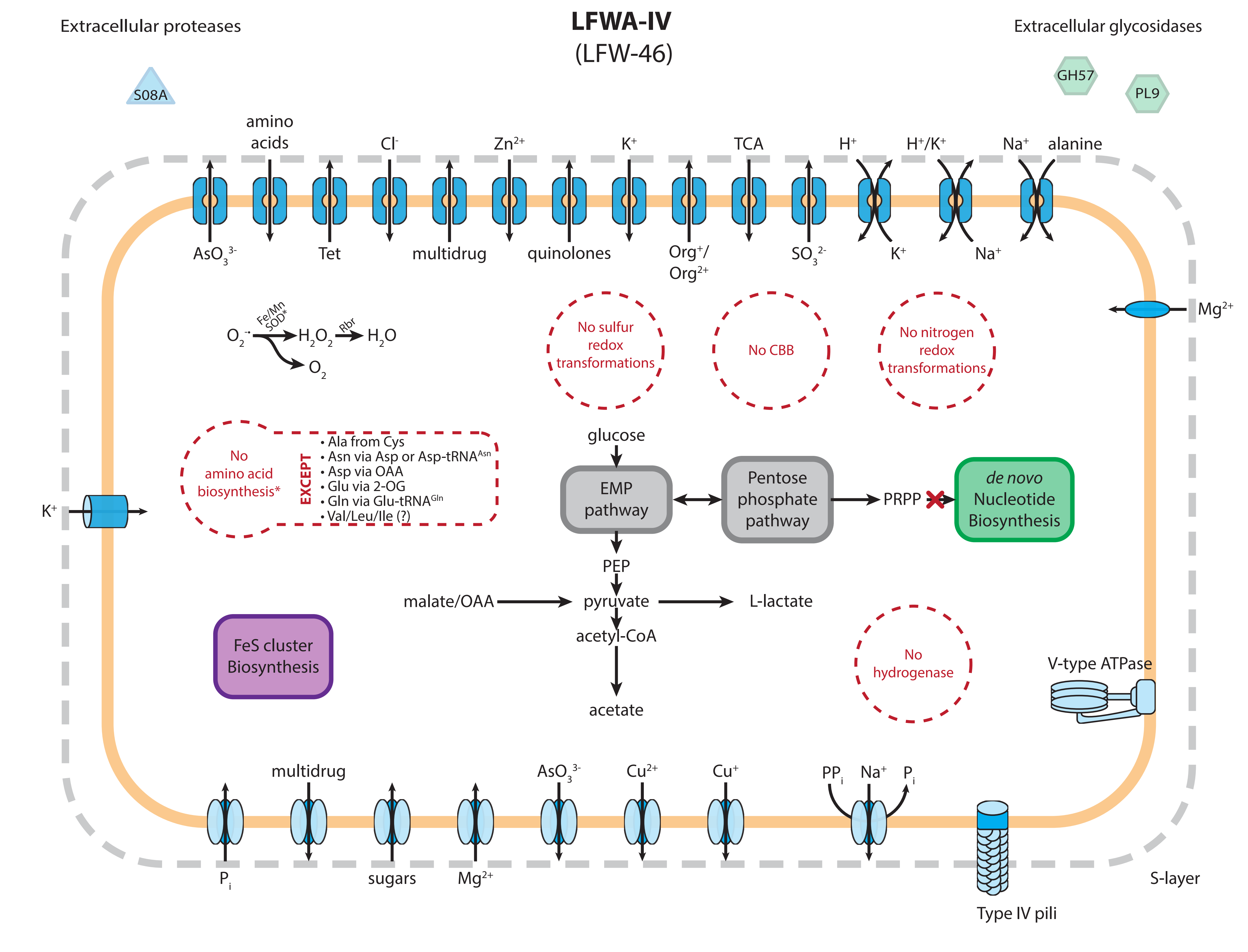

### Figure S5A

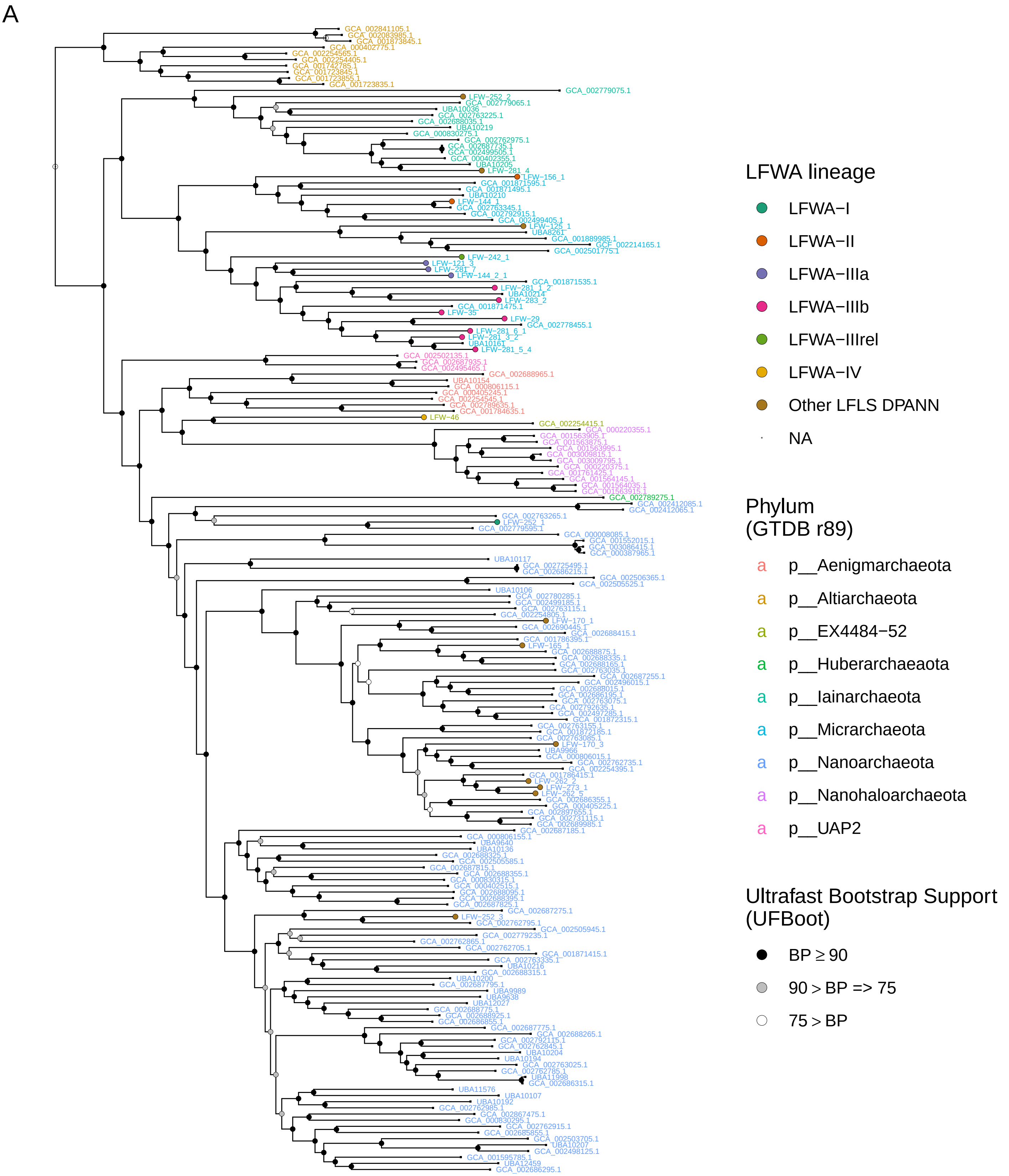

### Figure S5B

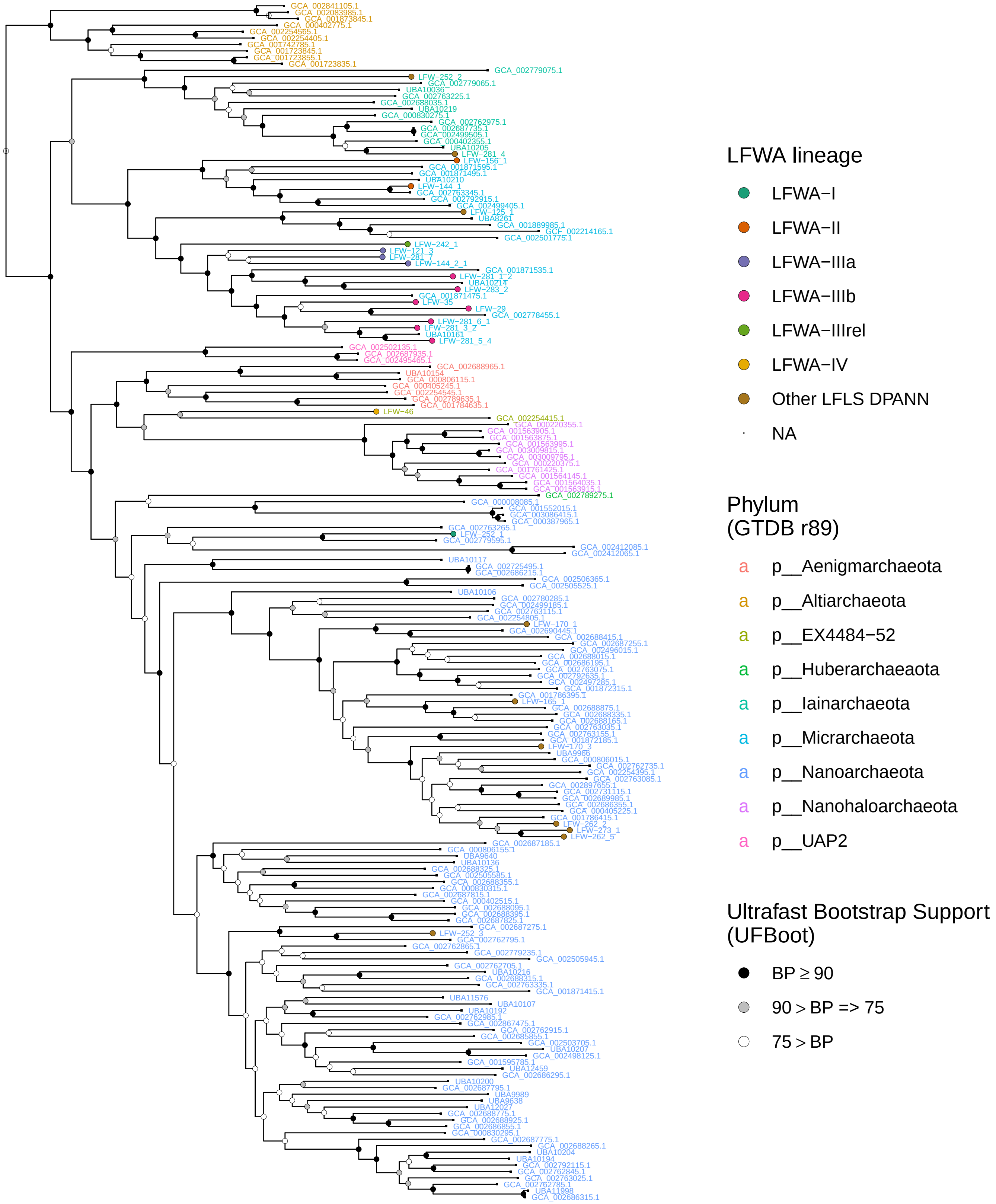

### Figure S5C

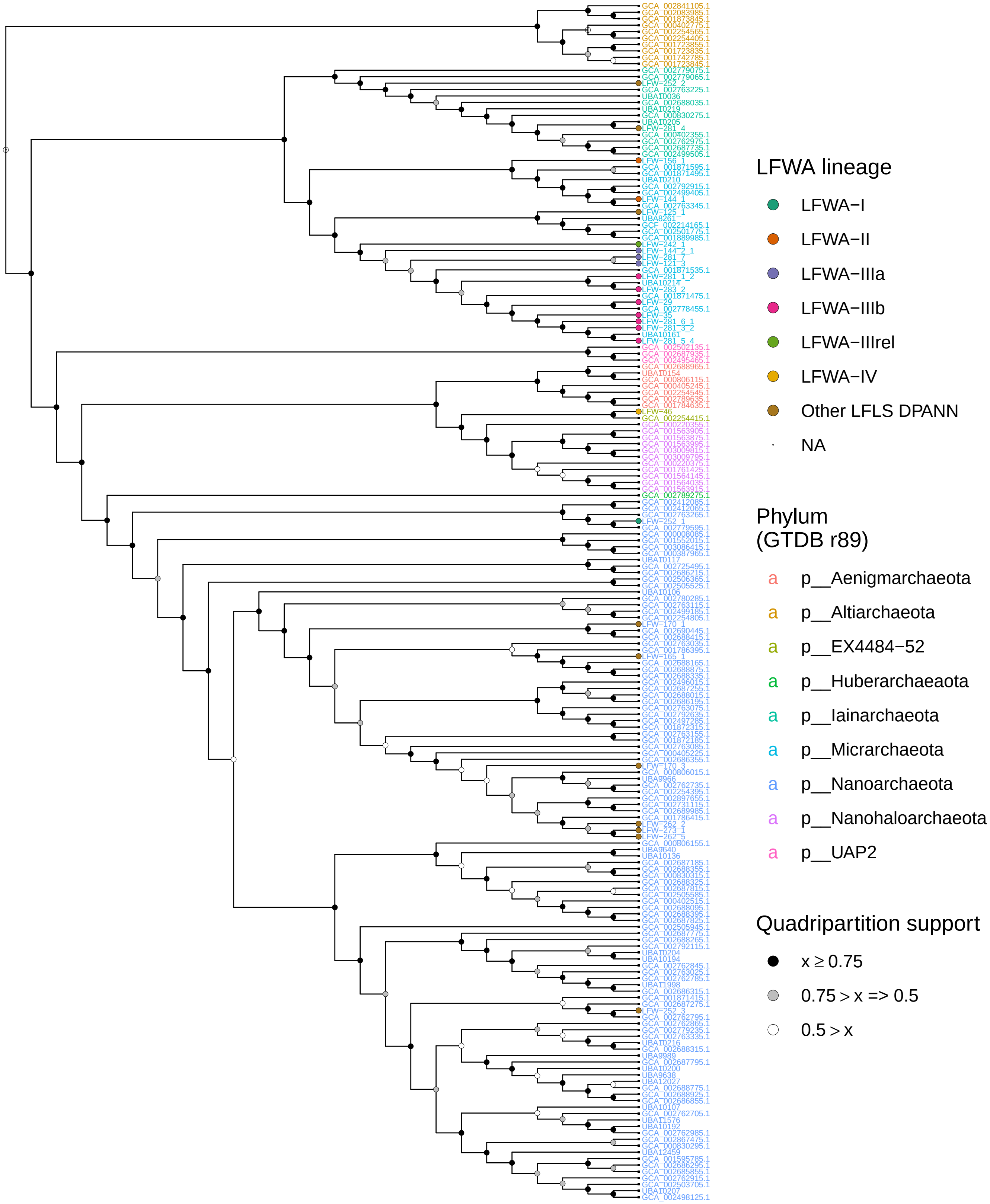

### Figure S6

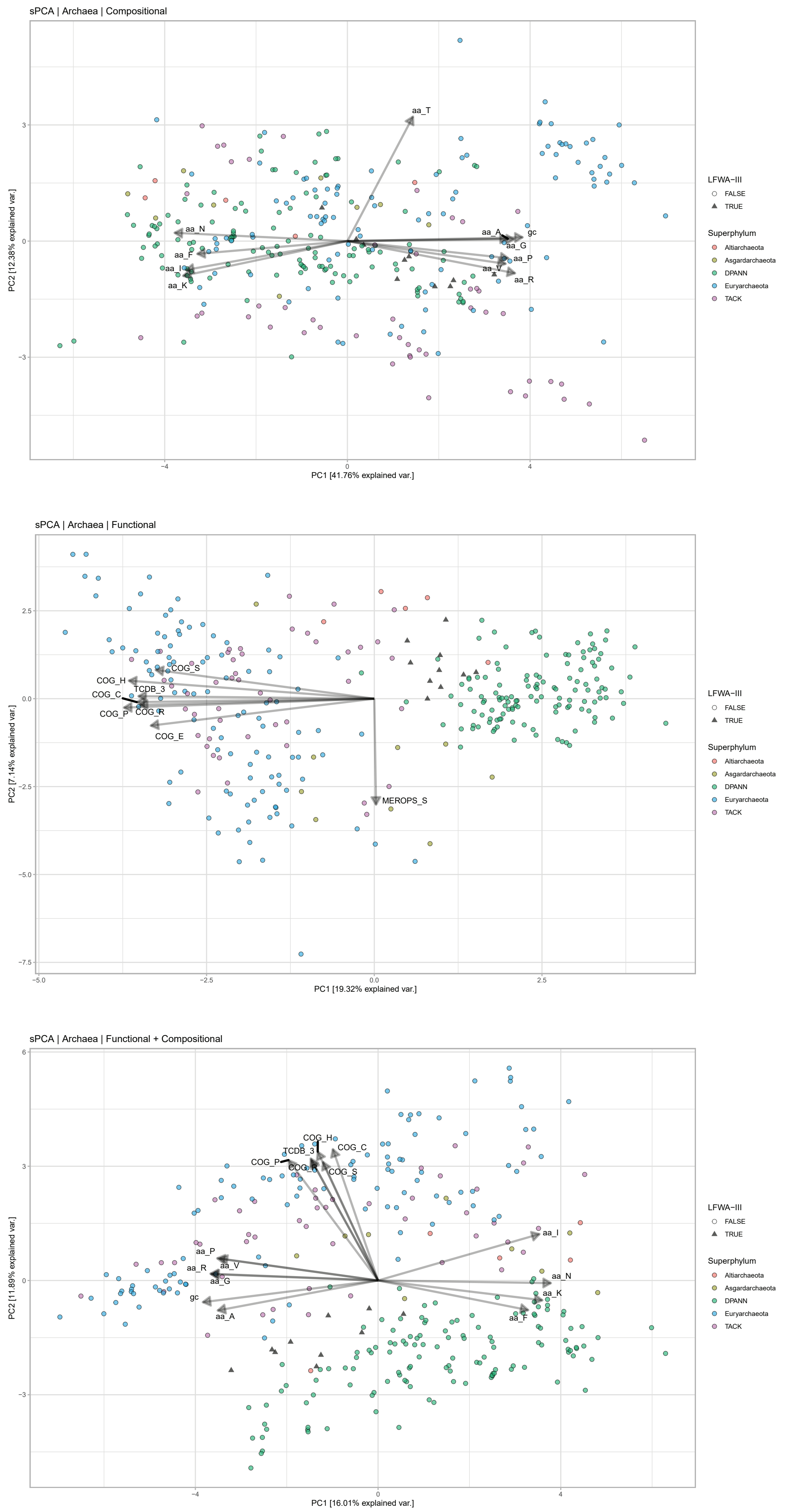

### Figure S7

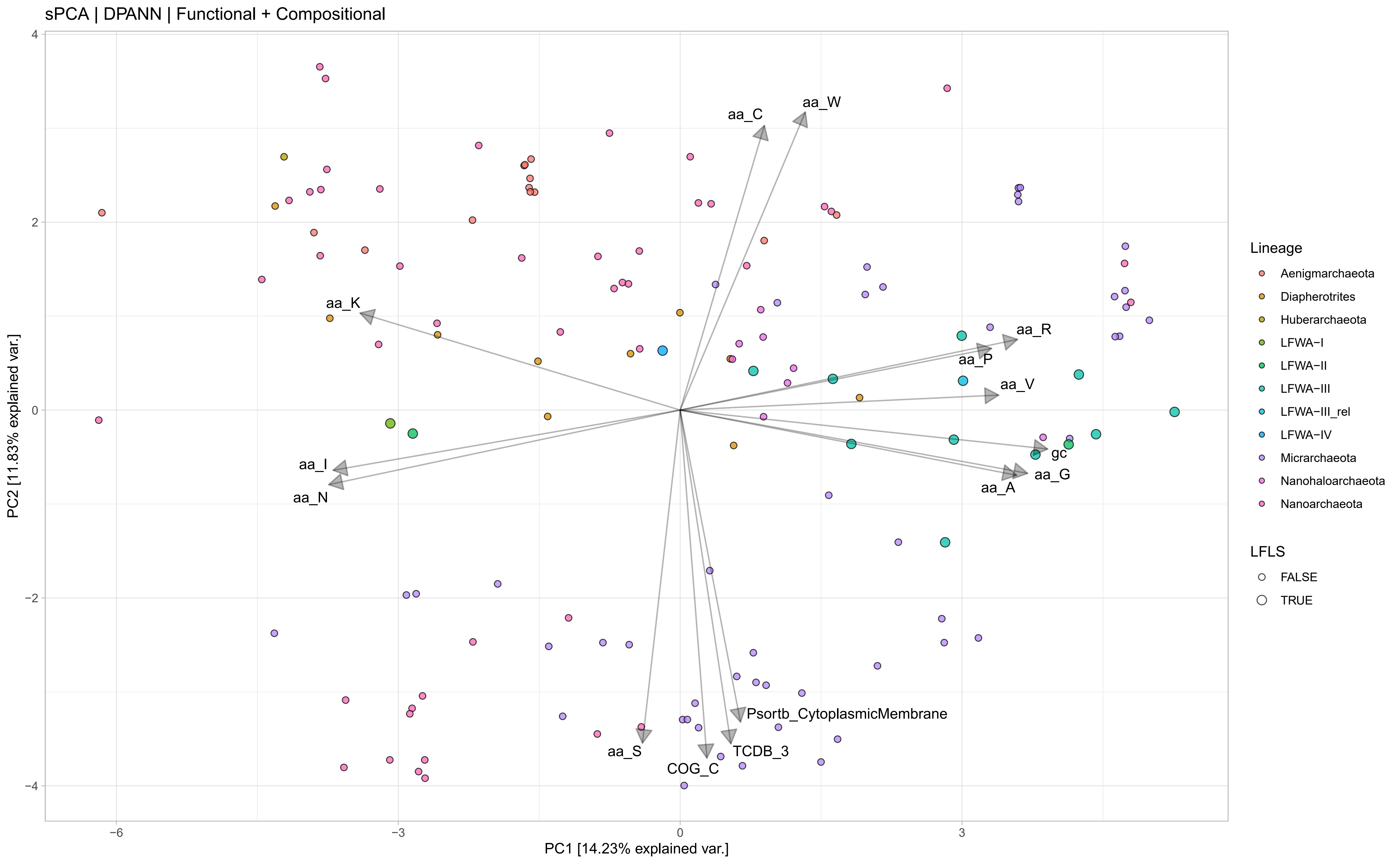

### Figure S8

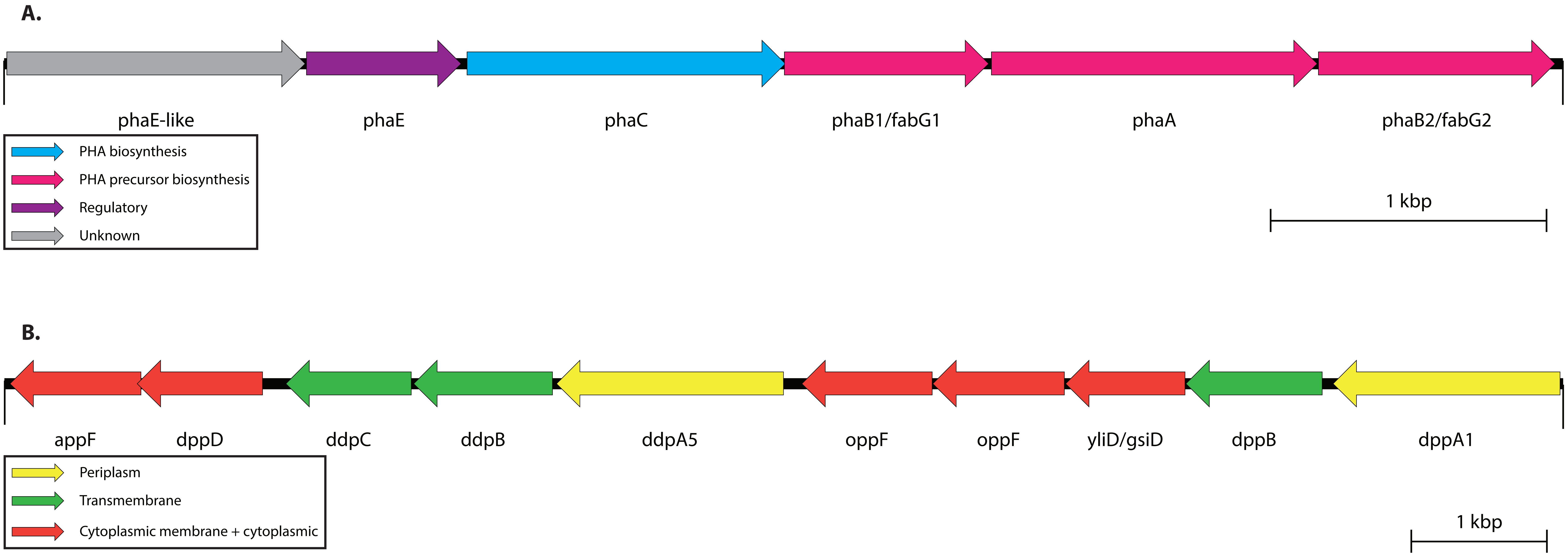

### Figure S9

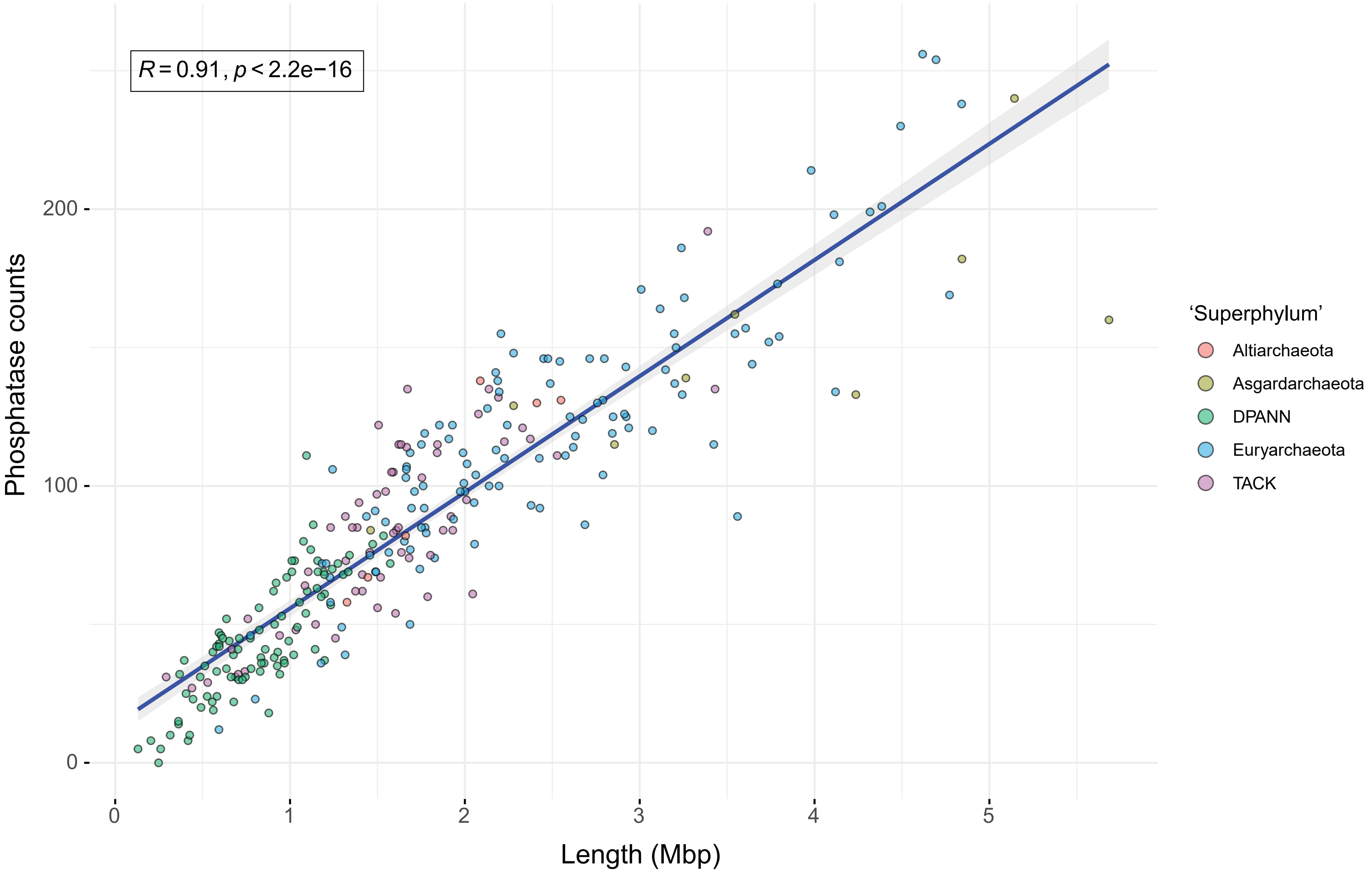

### Figure S10

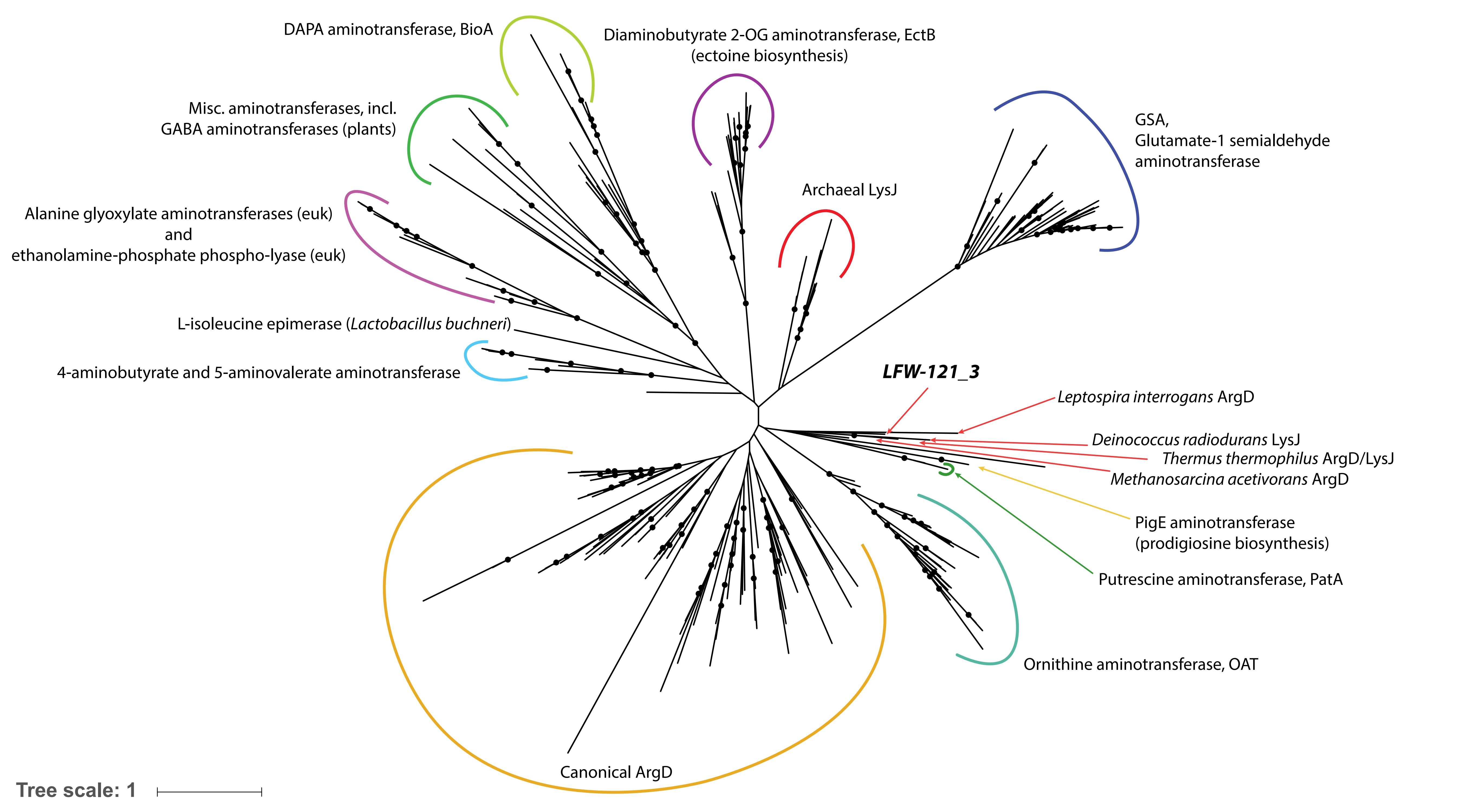
